# DNA polymerase diversity reveals multiple incursions of Polintons during nematode evolution

**DOI:** 10.1101/2023.08.22.554363

**Authors:** Dae-Eun Jeong, Sameer Sundrani, Richard Nelson Hall, Mart Krupovic, Eugene V. Koonin, Andrew Z. Fire

## Abstract

Polintons are dsDNA, virus-like self-synthesizing transposons widely found in eukaryotic genomes. Recent metagenomic discoveries of Polinton-like viruses are consistent with the hypothesis that Polintons invade eukaryotic host genomes through infectious viral particles. Nematode genomes contain multiple copies of Polintons and provide an opportunity to explore the natural distribution and evolution of Polintons during this process. We performed an extensive search of Polintons across nematode genomes, identifying multiple full-length Polinton copies in several species. We provide evidence of both ancient Polinton integrations and recent mobility in strains of the same nematode species. In addition to the major nematode Polinton family, we identified a group of Polintons that are overall closely related to the major family, but encode a distinct protein-primed B family DNA polymerase (pPolB) that is related to homologs from a different group of Polintons present outside of the *Nematoda*. Phylogenetic analyses on the pPolBs support the evolutionary scenarios in which these extrinsic pPolBs that seem to derive from Polinton families present in oomycetes and molluscs replaced the canonical pPolB in subsets of Polintons found in terrestrial and marine nematodes, respectively, suggesting inter-phylum horizontal gene transfers. The pPolBs of the terrestrial nematode and oomycete Polintons share a unique feature, an insertion of a HNH nuclease domain, whereas the pPolBs in the marine nematode Polintons share an insertion of a VSR nuclease domain with marine mollusc pPolBs. We hypothesize that horizontal gene transfer occurs among Polintons from widely different but cohabiting hosts.

## Background

Polintons/Mavericks (hereafter referred to as Polintons) were discovered as double-stranded DNA (dsDNA) transposons that encode a self-synthesizing, protein-primed DNA polymerase of the B family (pPolB) and a retroviral-element-like integrase (INT) (hence the name Polintons)^1,2^. Thus far primarily identified and characterized *in silico*, Polintons are among the larger known DNA transposons found widely across unicellular and multicellular eukaryotes, ranging from 13-25 kilobase pair (kbp) with 100-1500 base pair (bp) terminal inverted repeats (TIR) and with 5-8 bp target site duplications (TSD)^1-5^. In addition to pPolB and INT, Polintons typically encode a core set of conserved genes encoding homologs of dsDNA virus proteins involved in virion morphogenesis, such as adenovirus-like maturation protease (PRO), genome packaging ATPase, and major and minor capsid proteins (MCPs and mCPs, respectively)^5-11^. Polintons generally occupy a low fraction of their host genome; however, there are genomes with much higher occurrence, such as the excavate *Trichomonas vaginalis* where Polintons expanded to occupy more than 30% of the genome^3,12-18^.

While Polintons were first thought to be transposable elements, recent experimental and metagenomic discoveries of Polinton-like virophages and Polinton-like viruses suggest that Polintons found in multicellular eukaryotic genomes are capable of transmission between host genomes as infectious viral particles^19-25^. A combination of inferred protein characteristics (the presence of genes encoding homologs of the major and minor capsid protein and the ATPase and protease involved in capsid maturation), strongly supports this possibility^5,7^. A dual capability as viruses and transposons^5,7,18^ would afford Polintons an ability to propagate both within and between genomes.

Nematode genomes provide an excellent opportunity to observe transmissible genetic elements^26,27^ from diverse environments. The nematode phylum includes both parasitic and free-living species, with the latter having successfully adapted to diverse environmental niches including soil, rotten fruits, freshwater and seawater^28-33^. Because nematodes interact with diverse types of organisms (viruses, bacteria, protists, animals, and plants)^34-38^, the phylum is likely to have been exposed to a broad cross section of potentially infectious or transmissible elements. Thus, nematode genomes serve as a potentially-broad detector for elements capable of interspecies transfer and acquisition. Although several RNA viruses have been reported to infect or to be associated with nematodes^35,39-46^, only limited information about possible dsDNA virus infection and evolution in nematodes is available^47,48^.

*In silico* analyses have identified a handful of Polintons from nematode species^1,2,25,49^ including *C. briggsae*. These nematode Polintons encode a core set of Polinton proteins including pPolB, INT, ATPase, PRO, mCP and MCP, and nematode Polinton-specific protein PC1^1^, which was recently proposed to function as a fusogen^50^. The presence of a fusogen suggests that nematode Polintons may form an enveloped viral particle^51^. However, the natural distribution and evolution of Polintons across nematode genomes have not been extensively studied.

In this study, we use the diversity of available nematode genome sequences to investigate the natural variation and history of Polintons in this phylum. Of particular note, we observed an apparently modular history of the elements, with two distinct classes of pPolBs in otherwise-similar nematode Polintons. Additional analysis of these two classes outside of *Nematoda* suggests that Polintons have acquired and exchanged pPolB genes during evolution. The distribution of pPolB homologs in the Earth biome suggests that Polintons act as gene transfer vehicles.

## Results and Discussion

### Two groups of Polintons encoding distinct DNA polymerases in *Caenorhabditis* nematodes

To explore the distribution and evolution of Polintons in nematodes, we searched for full-length Polintons across 354 nematode genomes from 180 species and identified 266 Polintons in 66 genomes of 29 species (Extended Data Table 1; see Methods). While surveying the identified Polinton sequences, we noticed that a group of nematode Polintons encompassed a pPolB that was only weakly similar to the typical pPolB encoded by other nematode Polintons, in contrast to all other core Polinton genes and terminal inverted repeats (TIRs) that were highly conserved. We denoted the distant pPolB group ‘pPolB2’ to distinguish it from the typical nematode Polinton pPolBs (hereafter ‘pPolB1’) (Fig 1A).

**Fig 1.**
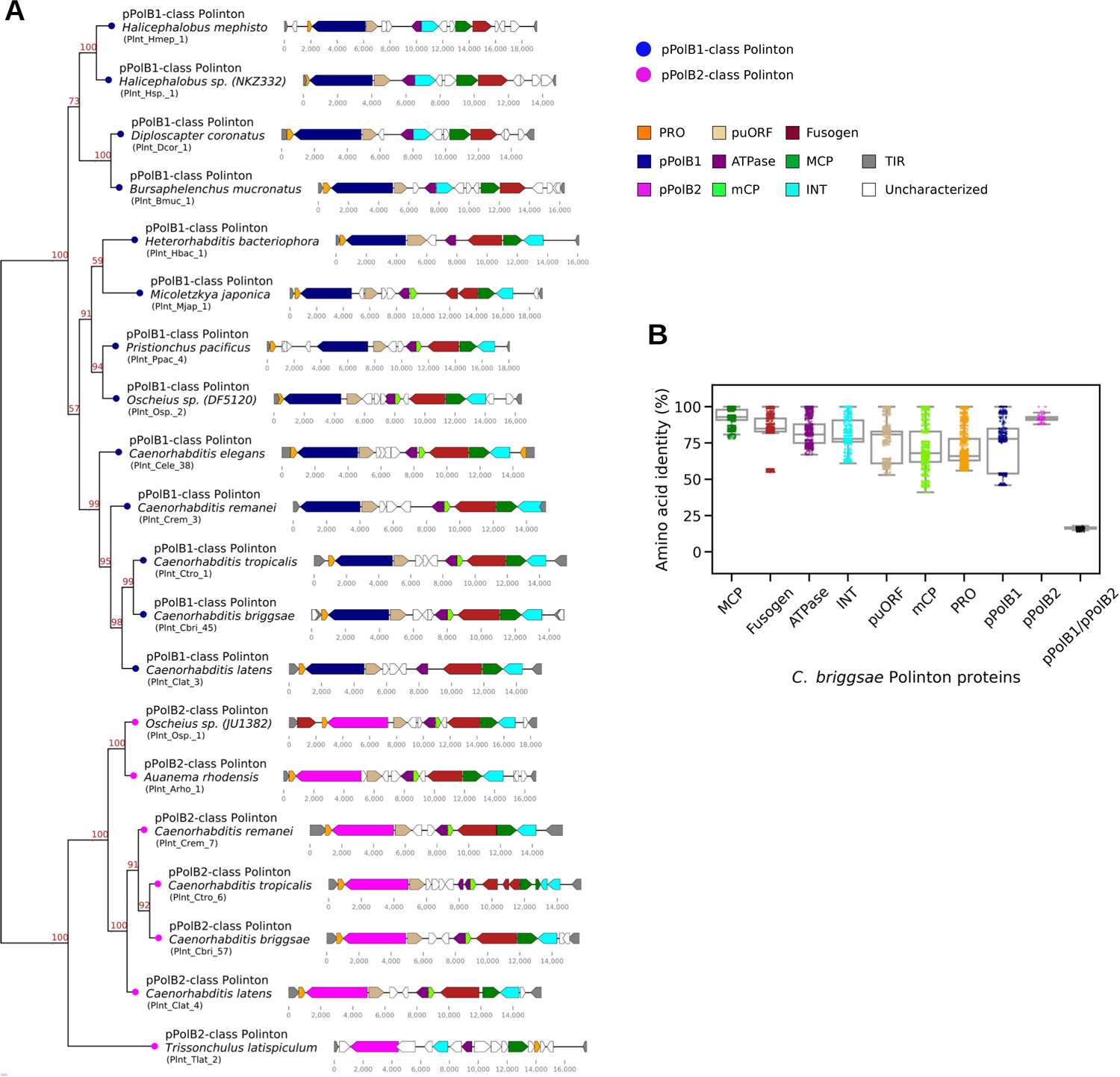
Two distinct DNA polymerase B families in *Caenorhabditis* Polintons. **A.** A phylogenetic tree (left) was constructed by applying a maximum likelihood-based method (IQ-TREE) to a multiple sequence alignment of pPolB proteins from Polintons of 19 nematode species. The genetic architectures of these elements are shown on the right. These Polintons were chosen for initial sequence alignment based on uninterrupted DNA polymerase and other Polinton protein coding regions. Navy and magenta colors of tree nodes indicate pPolB-1- and pPolB2-class Polintons, respectively. Bootstrap supporting values are shown on the top position of tree branches. Colors in right panels denote conserved genes encoding Polinton proteins; orange for an adenovirus-like maturation protease (PRO), navy and magenta for protein-primed DNA polymerase B genes (pPolB1 and pPolB2, respectively), beige for Polinton-uncharacterized ORF (puORF), purple for a packaging ATPase, lawngreen and cyan for major and minor capsid proteins (MCP and mCP, respectively), dark red for a fusogen, green for a retorviral-like-element integrase (INT), and white for novel (potentially either spurious or functional) ORFs that have not been previously described in other Polintons. Labels in parentheses denote unique identifiers (UI) for each Polinton (see Extended Data Table 1). **B**. A dot plot and a background box plot show percent (%) amino acid identities between each Polinton protein found in 49 *C. briggsae* Polintons.

The unexpected identification of two distinct groups of pPolBs in the nematode Polintons prompted us to examine the molecular diversification of Polinton proteins in nematodes in greater detail. To compare the extent of protein sequence similarity among different Polinton proteins, we constructed the corresponding multiple sequence alignments and calculated the percentage amino acid identity for each pair of homologous proteins from Polintons of *C. briggsae* (*C. briggsae* was chosen because this species was found to encompass the greatest number of both pPolB1- and pPolB2-class Polintons). The resulting comparisons showed 92% median identity for MCP, 85% for Fusogen, 81% for ATPase, 80% for INT, 66% for PRO, 78% for puORF, 68% for mCP, 77% for pPolB1 and 91% for pPolB2 (Fig 1B), indicating that these proteins are highly conserved among *C. briggsae* Polintons.

In a sharp contrast, pairwise comparisons between pPolB1 and pPolB2 showed a median of 16% amino acid sequence identity. The Polinton pPolBs belong the protein-primed family B DNA polymerases that contain conserved exonuclease motifs (Exo I, II, and III) required for proofreading activity and polymerase motifs (Pol A, B, and C) required for DNA polymerization activity^52-55^. Examination of the multiple sequence alignment of representative pPolB1 and pPolB2 proteins from Polintons of 19 nematode species with the prototypical Phi29 phage pPolB indicated that all three exonuclease and all three polymerase motifs are conserved in both pPolB1 and pPolB2 (Extended Data Fig 1 and Extended Data Fig 2A), suggesting that both groups of the nematode Polintons possess fully functional DNA polymerases.

Given the low sequence similarity between pPolB1 and pPolB2, we also compared the structural models of the two proteins obtained using AlphaFold2^56^ through ColabFold^57^. Structural alignment and superposition of *C. briggsae* pPolB1 and pPolB2 structures show evidence of a close structural similarity between the two groups of pPolBs, in spite of the low sequence identity, in line with the prediction that both are active DNA polymerases (Extended Data Fig 2B-D and Fig 2A). In addition to the conserved structural core of pPolB1 and pPolB2, the structural alignment also revealed potential pPolB1- and pPolB2-specific domains (Fig 2A). In particular, we identified a pPolB2-specific domain that is conserved across all nematode pPolB2 proteins except the pPolB2 from three Polintons in a marine species, *Trissonchulus latispiculum* (Fig 2A and Extended Data Fig 1). A protein domain search using HHpred^58^ identified this domain as a putative HNH endonuclease^59^ (Fig 2B-E). An additional structural alignment between terrestrial *C. briggsae* pPolB2 and marine *T. latispiculum* (*Tlat*) pPolB2 revealed a *Tlat* pPolB2-specific domain, which was subsequently identified as a VSR endonuclease domain (Fig 2G-J). Apparently, this pPolB2 family acquired an additional endonuclease domain during its evolution.

**Fig 2.**
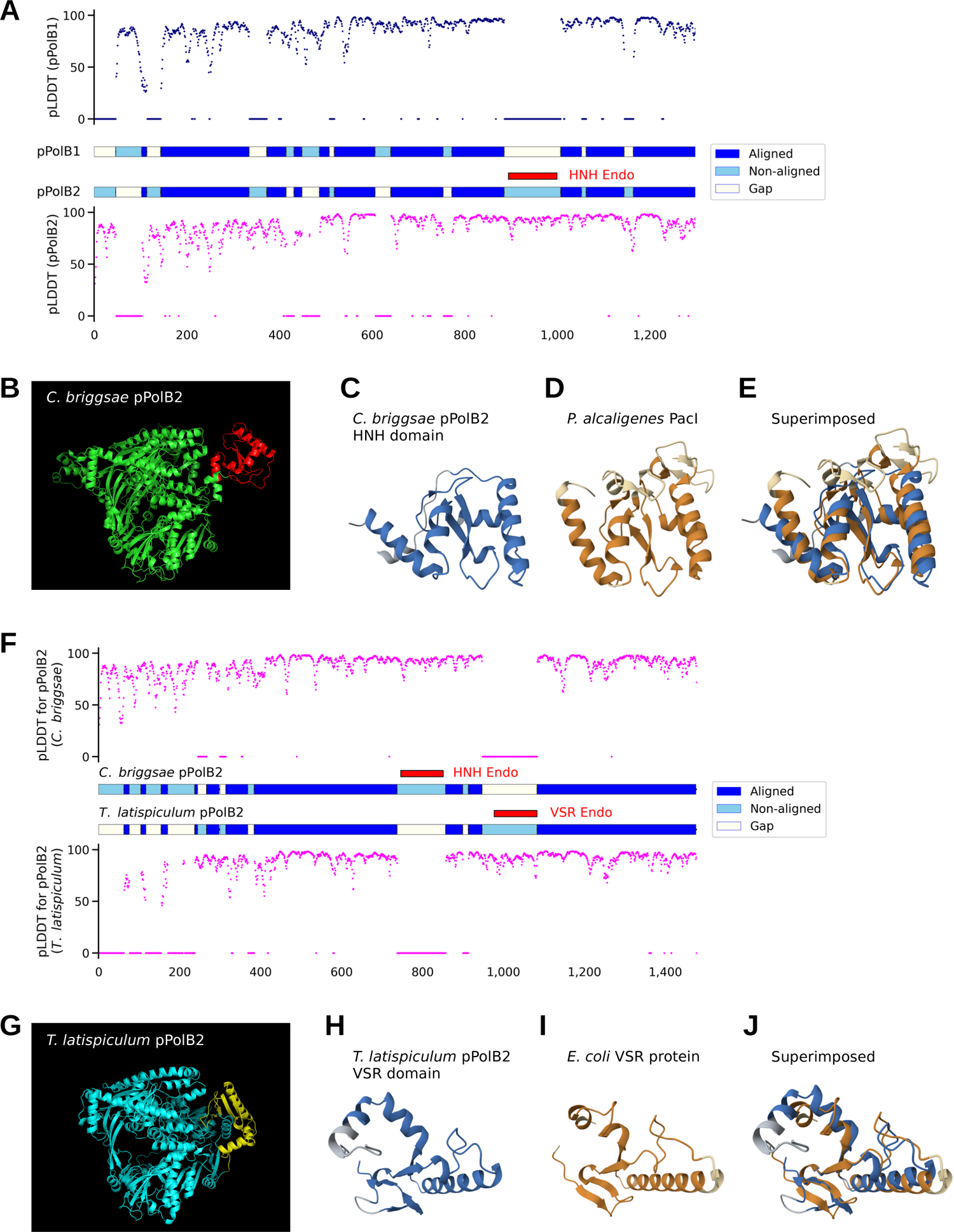
HNH and VSR endonuclease domains are found in terrestrial and marine nematode pPolB2s, respectively, but not in pPolB1. **A.** Aligned (blue), non-aligned (skyblue) and gap (white) parts of pPolB1 and pPolB2 (middle panel) from sequence-free structural alignment are shown with pLDDT (predicted local distance difference test) score predicted at the single amino acid level by AlphaFold2. Overall aligned parts for both pPolB1 and pPolB2 showed high pLDDT score, suggesting confident prediction of the structures. **B.** The predicted structure of *C. briggsae* pPolB2 (green) with the HNH endonuclease domain (red). **C-E.** The predicted structure of pPolB2-specific HNH domain (blue, **C**) and the crystal structure of *P. alcaligenes* PacI HNH endonuclease (orange, **D**) (PDBid: 3M7K)^59^ are superimposed (**E**) (RMSD: 3.03, TM-score: 0.55) to show potentially conserved structural folds. **F.** Aligned (blue) and non-aligned (skyblue) parts of *C. briggsae* pPolB2 and *T. latispiculum* pPolB2 (middle panel) from sequence-free structural alignment are shown with pLDDT score. **G.** The predicted of *T. latispiculum* pPolB2 (cyan) with the VSR domain (yellow). **H-J.** The predicted structure of *T. latispiculum* pPolB2 VSR endonuclease domain (blue, **H**) and the crystal structure of *E. coli* VSR endonuclease (orange, **I**) (PDBid: 1VSR)^85^ are superimposed (**J**) (RMSD: 3.06, TM-score: 0.63) to show potentially conserved structural folds. Dark and light colors for both orange and blue (**C-E** and **H-J**) indicate aligned and non-aligned positions, respectively.

Taken together, these observations suggest an ancient separation between pPolB1 and pPolB2 polymerase families. In the following sections, we investigate the potential origins and evolution of the two groups of nematode Polintons.

### Inter- and intra-species copy number variations of pPolB1 and pPolB2 classes of Polintons in nematode genomes

An extensive search for Polintons in diverse nematode genomes revealed variations in the inter- and intra-species distributions and copy numbers of the pPolB1 and pPolB2 groups of Polintons. The pPolB1-class and pPolB2-class Polintons were detected in 25 and 12 nematode species, respectively (out of a total of 180 species) (Fig 3 and Extended Data Table 1).

**Fig 3.**
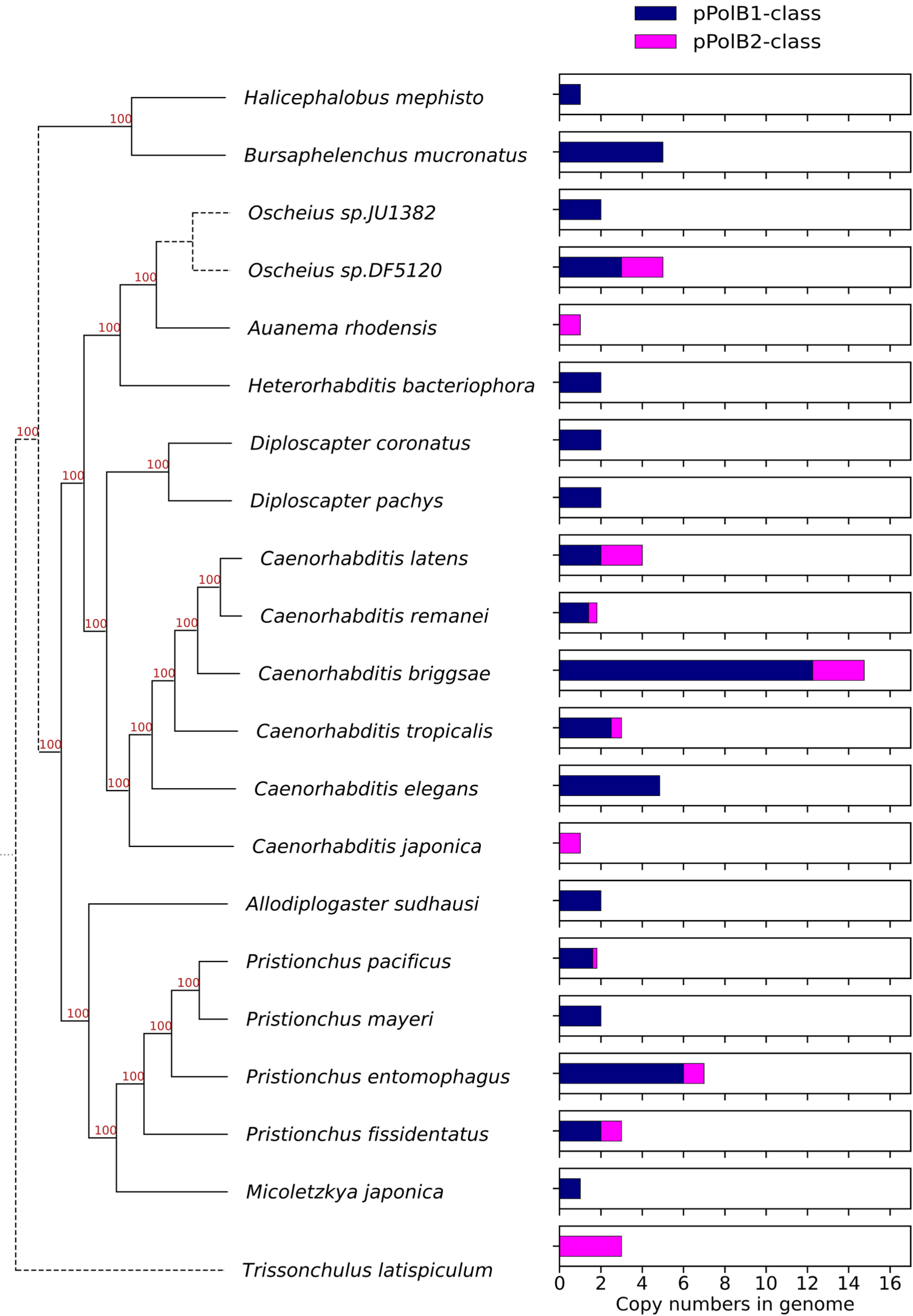
Inter-species copy number variations of nematode Polintons. Bar graphs (right panel) showing average copy numbers per genome of full-length pPolB1-class (navy) and pPolB2-class (magenta) Polintons in *Nematoda* species are shown with a nematode species phylogenetic tree (left panel) that was obtained from Ahmed et al. (2022)^86^.

Polintons of both classes were detected in several nematode genera (*Pristionchus*, *Oscheius*, and *Caenorhabditis*), with varying copy numbers; notable copy number variations were also observed between species within a genus, in particular, *Caenorhabditis*, for which multiple genomes are available (Fig 3). Generally, the pPolB1-class Polintons were substantially more abundant than the pPolB2-class ones, with several species carrying only pPolB1 Polintons and only four possessing pPolB2 Polintons (Fig 3 and Extended Data Table 1). These observations appear to be compatible with independent spread of the pPolB1 and pPolB2 classes of Polintons among nematodes.

We also observed distinct patterns and copy numbers of Polintons within each species, most clearly, in a set of *Caenorhabditis* species for which genome sequences are available for multiple wild-type strains (Extended Data Fig 3A-D). Because genomes for three wild-type strains of *C. briggsae* have been assembled at chromosome level^60,61^ with the largest copy numbers of both pPoB1- and pPolB2-class Polintons in nematodes (Fig 3 and Extended Data Fig 3D), we further analyzed the genomic neighborhoods of the Polintons, to test whether transposition occurred before or after the diversification of the wild-isolates. Only two pPolB1-class Polintons shared the same positions in all three *C. briggsae* strains and 6 pPolB1-class Polintons shared the positions in AF16 and QX1410 strains (Extended Data Fig 3E), suggesting that these Polintons already occupied the same positions in the genomes of the respective *C. briggsae* ancestors. In contrast, the remaining pPolB1-class Polintons and all pPolB2-class Polintons were found at unique positions (Extended Data Fig 3E), suggesting transposition events that occurred after the strain divergence. These intra-species variations imply that Polintons were actively integrating into *Caenorhabditis* genomes both before and after the diversification of the characterized strains, with each genome reflecting a distinct natural history of exposure to the two classes of Polintons.

### Transfer of pPolB genes among Polintons

The discovery of two distinct groups of pPolBs in nematode Polintons could be explained through one of two alternative scenarios. Under the first scenario, there are two distinct classes of nematode Polintons with only an ancient relationship, in which case, all proteins would show low sequence conservation, in the same range as the pPolBs. The second scenario involves a relatively recent pPolB swap, in which case, the other proteins would be far more similar than the pPolBs.

The bulk protein sequence comparisons described above favor the second scenario, but to investigate the evolution of the two groups of Polintons further, we examined the relationships among the Polintons within a single species (*C. briggsae*) by calculating sequence similarities for all possible pairwise combinations of Polintons. This analysis showed strong positive correlations between the pairwise similarities of all proteins other than pPolB, regardless of whether the compared Polintons encoded pPolB1 or pPolB2 (Fig 4A-B and Extended Data Fig 4A-B). In within-class comparisons, strong correlations between the similarities among pPolBs and those among other Polinton proteins (Fig 4C) suggest that pPolB1 and pPolB2 proteins can be maintained within Polintons through substantial evolutionary spans. By contrast, in between-class comparisons, the similarities among pPolB1 and pPolB2 did not correlate with those among other proteins. (Fig 4C). These observations strongly suggest that in one of the nematode Polinton classes pPolB was relatively recently replaced by a distantly related homolog.

**Fig 4.**
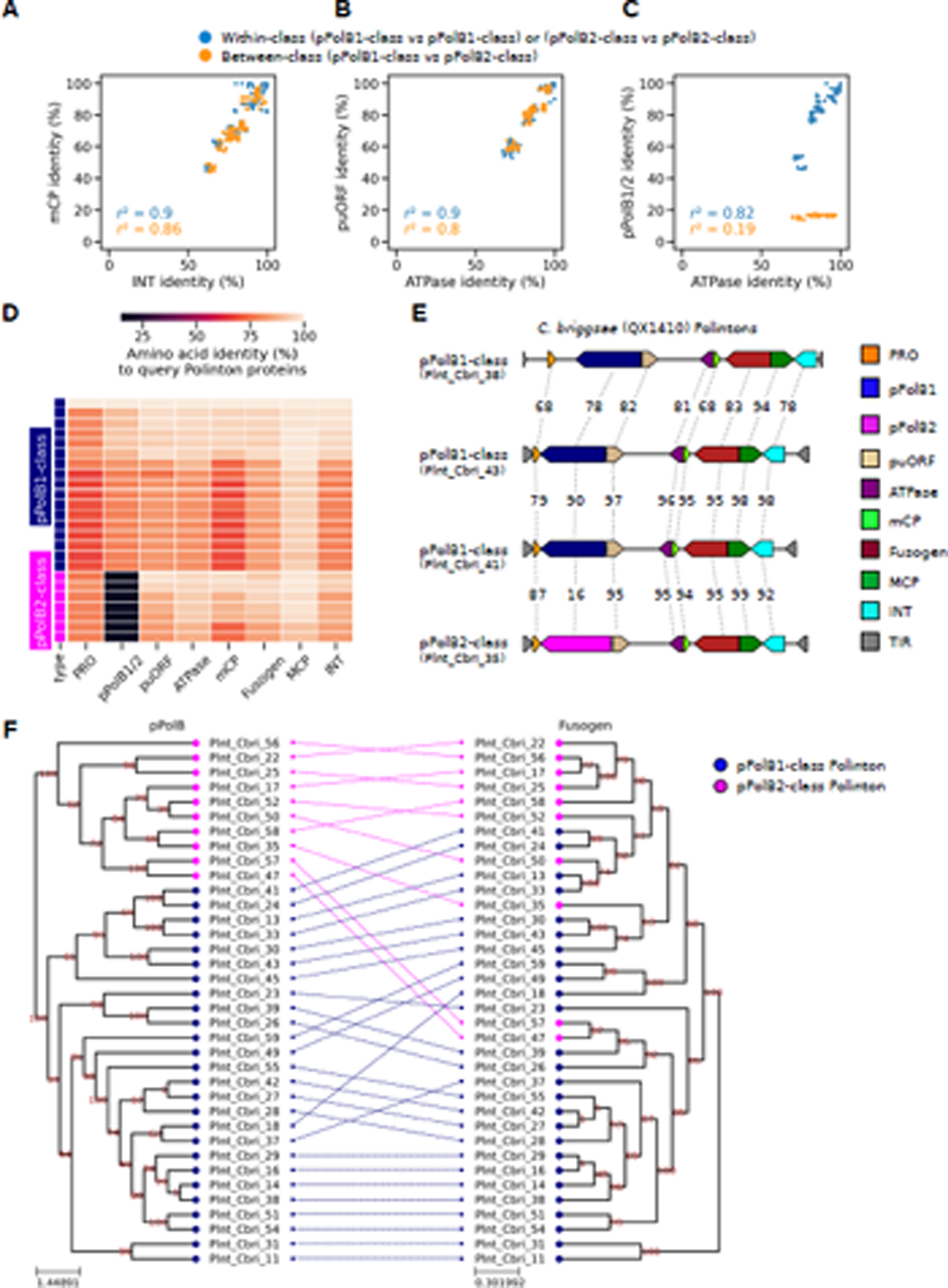
Two distantly related groups of pPolBs in *C. briggsae* Polintons. **A-C**. Scatter plots showing percent amino acid identity of one protein for each axis from *C. briggsae* Polinton species; mCP and INT (**A**), puORF and ATPase (**B**), and pPolB1/2 and ATPase (**C**). Each dot indicates a pair of two *C. briggsae* Polintons, whose proteins indicated at the X and Y axes were aligned, respectively, for calculating percent identity. Dot colors indicate whether two Polintons in a pair for comparison are within the same pPolB group (blue) or from different pPolB groups (orange). **D.** Heatmap showing percentage amino acid identity of proteins. Each row shows an individual *C. briggsae* Polinton (navy for pPolB1-class and magenta for pPolB2-class), while each column represents one of the conserved Polinton proteins. All comparisons were performed against a query *C. briggsae* Polinton (Plnt_Cbri_18). **E.** Genetic architectures of four *C. briggsae* Polintons (QX1410 strain, a wild isolate) shown with % amino acid identities for each Polinton protein. Plnt_Cbri_41 and Plnt_Cbri_43 are relatively close to each other, showing > 90% identity, while Plnt_Cbri_43 and Plnt_Cbri_38 are more distant (evidenced by lower % identities). The continuous nature of the homology outside of the pPolB region provides evidence for gradual divergence in these regions during the ancestry of the current elements. By contrast, the Plnt_Cbri_35 Polinton showed strong similarity in all proteins except pPolB (particularly to PInt_Cbri_41), with dramatic divergence for pPolB, suggesting a swapping event of the pPolB gene during the derivation of this Polinton. **F**. A tanglegram showing the relationship between two phylogenetic trees constructed from multiple sequence alignment of *C. briggsae* pPolB1/2 (left panel) and Fusogen (right panel) proteins. The colors of the nodes indicate pPolB1-class (navy) and pPolB2-class (magenta). Dashed lines between two trees denote the correspondence of pPolB and Fusogen proteins found from the same Polinton. The tree exemplifies that close relationships between non-pPolB proteins can be accompanied by very distant relationships between the corresponding pPolBs.

We further searched for examples of pPolB1 and pPolB2 genes being embedded in otherwise closely similar Polinton sequences. One such example, providing a useful illustration, was observed in *C. briggsae* strain QX1410 (Fig 4D-E), where Polintons Plnt_Cbri_41 and Plnt_Cbri_35 both showed high similarity to the canonical Plnt_Cbri_43 sequence in all non-pPolB proteins (Fig 4D), whereas the pPolBs showed dramatic divergence (only 16% sequence identity) (Fig 4E). A more general view of these relationships is apparent from the phylogenetic trees of the pPolBs and the rest of the Polinton proteins (Extended Data Fig 5A-H). In the pPolB tree, the two groups of pPolBs formed two well separated clades whereas the other Polinton proteins associated with either pPolB1 or pPolB2 were interspersed in the respective trees (Fig 4F, and Extended Data Fig 4C and 5A-H). In addition to *C. briggsae*, we also observed a possible inter-Polinton exchange of pPolB genes in an *Oscheius* species (DF5120) as indicated by the observation that an MCP protein from pPolB2-class Polinton clustered with a pPolB1-class Polinton MCP in the phylogenetic tree (Extended Data Fig 6A-B). These observations are compatible with the proposed exchange of distinct pPolB genes between Polintons.

### Distribution of pPolB1 and pPolB2 homologs across eukaryotes suggests distinct evolutionary trajectories of pPolB1 and pPolB2

To characterize the evolutionary trajectories of the two pPolB types on a larger evolutionary scale, we explored the distribution of pPolB1 and pPolB2 homologs across the available eukaryotic genome assemblies (7,897 assemblies from 64 phyla; 1.63 Terabases available through NCBI genome database as of late April, 2023). To focus on Polintons, we restricted our analysis to pPolB genes linked to INT homologs (see Methods). We identified 20,702 INT-associated pPolB1 and 1,331 pPolB2 homologs distributed across 960 and 164 genomes of diverse eukaryotes, respectively (Extended Data Table 2 and 3). These pPolBs found from all taxon groups contained conserved exonuclease and polymerase motifs for pPolB1 and pPolB2 groups (Extended Data Fig 7), suggesting that these pPolBs are functional Polinton polymerases.

Distinct patterns of pPolB1 and pPolB2 distribution at the phylum level were identified (Fig 5A). Mimicking the case of nematodes, pPolB1-containing Polintons showed a wide spread across 30 phyla. pPolB1 were found to be most abundant in invertebrates, with multiple copies found in 17 of the 23 invertebrate phyla. By contrast, pPolB2 showed a sparse distribution, being represented mostly in two unicellular eukaryote phyla, *Metamonada* and *Oomycota*, and five invertebrate phyla, *Annelida*, *Brachiopoda*, *Echinodermata*, *Mollusca*, and *Nematoda*. The highest pPolB2 copy numbers were detected in *Metamonada*, particularly, in *Trichomonas vaginalis* (388 copies) and *Tritrichomonas foetus* (259 copies), consistent with a previous report on the high abundance of Polintons in *T. vaginalis*^3^.

**Fig 5.**
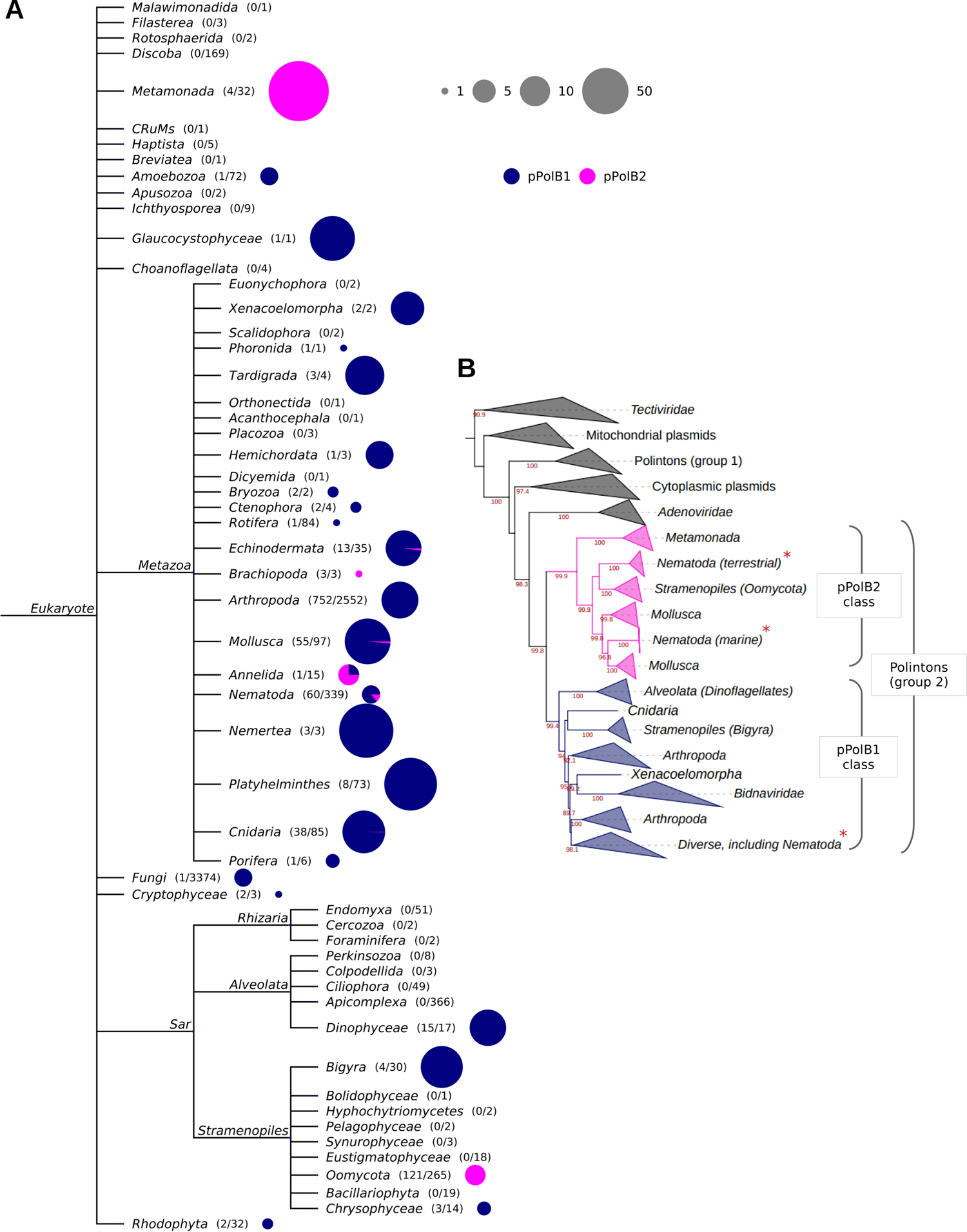
Distinct distribution and evolutionary trajectories of pPolB1 and pPolB2 groups in the biosphere. **A.** A NCBI taxonomy tree with Eukaryotic taxa. The numbers in parentheses indicate the number of genomes having at least one copy of pPolB and the number of total genomes subjected to the Polinton pPolB search (see Methods for details). The sizes and colors in the pie charts indicate the copy number of Polinton pPolBs per pPolB-containing genomes and the classes of pPolB; navy for pPolB1 and magenta for pPolB2, respectively. **B.** A phylogenetic tree was constructed from multiple sequence alignment of pPolB proteins found in diverse taxa including viruses of prokaryotes, eukaryotes, DNA plasmids, and Polintons using the IQ-TREE-inferred maximum likelihood method. Bootstrap supporting values are shown on the bottom positions of tree branches. All pPolBs identified in this study were found within Group 2 Polinton pPolBs where two big monophyletic groups for pPolB1 and pPolB2 classes are shown with green and blue background colors, respectively. Red asterisks indicated the *Nematoda* pPolB1 and pPolB2 proteins identified in this study.

A broader phylogenetic analysis that included a set of representatives from the main groups of pPolBs encoded by viruses of prokaryotes and eukaryotes as well as Polintons and DNA plasmids further illuminated the evolutionary trajectories of pPolB1 and pPolB2 (Fig 5B). Each of the two groups of pPolBs was found to be monophyletic (Fig 5B and Extended Data Fig 8). pPolB1 and pPolB2 formed distinct clades in the tree, and their relative positions suggested that these two DNA polymerase forms diverged early on. Given the greater diversity and broader distribution among the nematodes of pPolB1 compared to pPolB2 (Fig 3 and Extended Data Fig 3A-E), it appears most likely that the ancestral nematode Polintons encoded pPolB1 which was replaced in one nematode branch. That replacement might have occurred via horizontal gene transfer from oomycete Polintons, given the tree topology and the highest sequence similarities between oomycete pPolBs and nematode pPolB2s (Fig 5B and Extended Data Fig 8). Furthermore, protein domain search showed that pPolB2 of oomycete Polintons shares the HNH endonuclease domain with the homolog from the nematode Polintons (Fig 6, and Extended Data Fig 9 and 10A), to the exclusion of other pPolB2s. Thus, the HNH domain is a unique synapomorphy (shared derived character) that links the oomycete and nematode Polintons encoding pPolB2, strongly supporting the horizontal gene transfer scenario. The HNH domain could potentially have been captured by the oomycete pPolB2 from a Group I self-splicing intron, before finding its way into the nematode genomes^62^.

**Fig 6.**
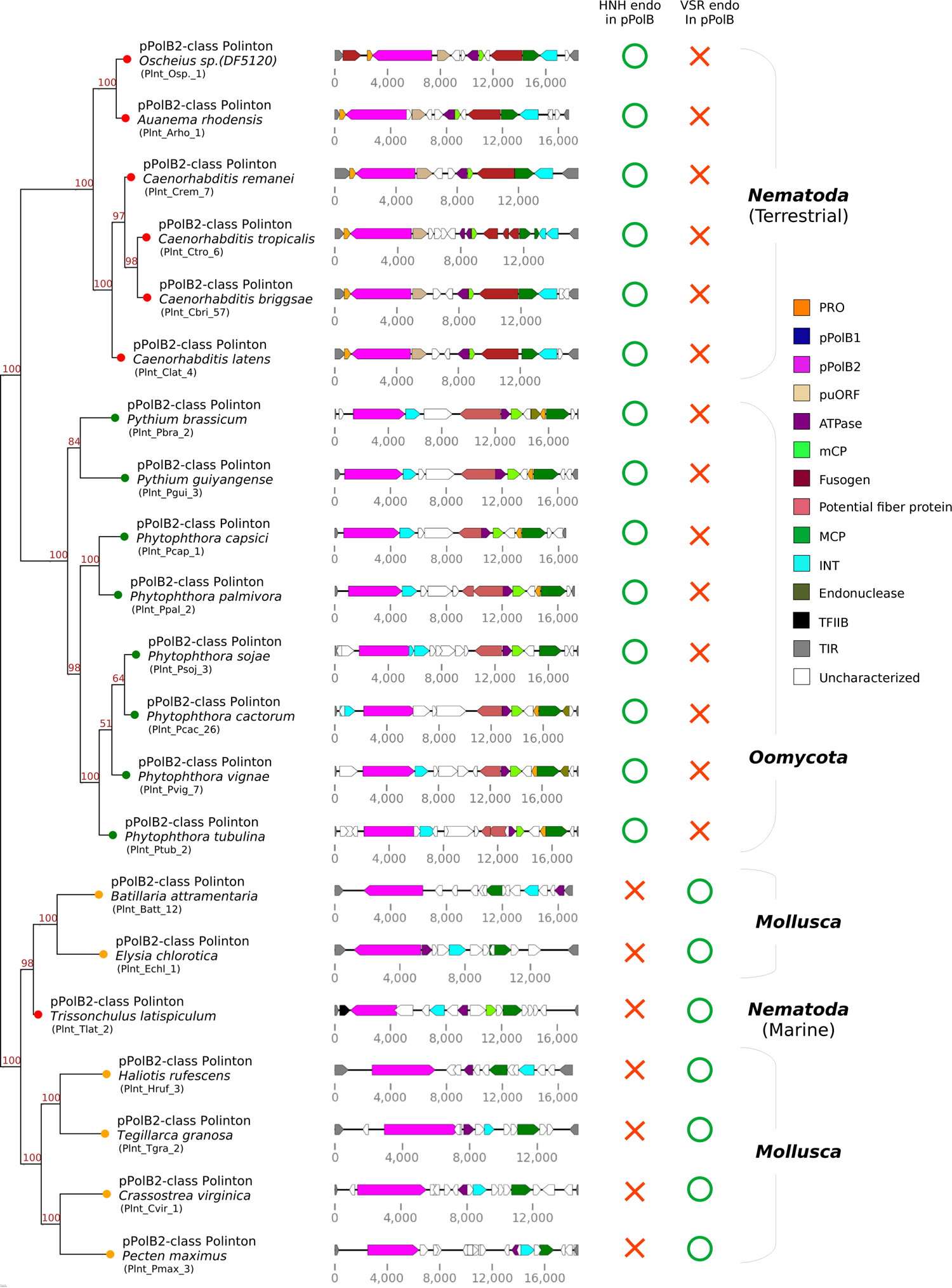
Phylogenetic relationships of Polinton pPolB2 proteins from three distinct phyla. A Phylogenetic tree (left) has been built by using IQ-TREE-inferred maximum likelihood on a multiple sequence alignment of pPolB2 protein sequences found from *Nematoda*, *Oomycota* and *Mollusca* (See Extended Data Fig 9 for full alignment data.). Bootstrap supporting values are shown on the top position of tree branches. Colors in right panels denote conserved genes encoding Polinton proteins; orange for an adenovirus-like maturation protease (PRO), navy and magenta for protein-primed DNA polymerase B genes (pPolB1 and pPolB2, respectively), beige for Polinton-uncharacterized ORF (puORF), purple for a packaging ATPase, lawngreen and cyan for major and minor capsid proteins (MCP and mCP, respectively), dark red for a fusogen, green for a retorviral-like-element integrase (INT), olive for an endonuclease, ivory for potential fiber protein and white for unknown ORFs.

While analyzing protein profiles of oomycete Polintons, we noted that oomycete Polintons encode a conserved protein showing similarity to beta-helix-fold tail fiber proteins of tailed bacteriophages (Fig 6), raising a possibility that these proteins mediate recognition and binding of oomycetes Polintons and oomycetes hosts. Notably, unlike nematode Polintons, oomycete Polintons do not encode membrane fusion proteins and hence are unlikely to be enveloped. Whether potential envelopment is an exclusive property of the nematode Polintons compared to Polintons from other organisms remains to be investigated.

Given the apparent relatively recent incursion of pPolB2 into nematode Polintons, direct inter-phyla horizontal transfer of the pPolB2 gene between oomycete and nematode Polintons, perhaps, via virus particles formed by Polintons appears likely. Under this scenario, a nematode would have been infected by the oomycete Polinton virus particles, with subsequent recombination with a resident nematode Polinton such that only the pPolB gene was transferred. Alternatively, the immediate source of the pPolB2 homologs in nematodes might have been an intermediate species. In either case, the pPolB2-encoding Polinton family was fixed in the nematodes and spread to a variety of nematode species, again, possibly, through Polinton virions.

### A distinct group of pPolB2 Polintons in a marine nematode species

A recent study of marine nematode genomes^63^ allowed us to further examine Polintons in an ecological niche for which genome data were initially limited. In addition to land-dwelling nematodes (both free and parasitic), there are many nematode species that dwell in aquatic habitats^28,32,33^. Notably, analysis of the recently sequenced marine nematode genomes identified a Polinton-containing genome (*Trissonchulus latispiculum*^63^) that encompasses no pPolB1-class Polintons but carries a distinct group of three pPolB2-class Polintons (Fig 1A and Extended Data Table 1). Sequence and structural comparison of the terrestrial *C. briggsae* pPolB2 and a marine *T. latispiculum* pPolB2 showed that *T. latispiculum* pPolB2 lacked the HNH nuclease domain but instead contained a VSR (very short patch repair) endonuclease domain (Fig 2F-G and Extended Data Fig 9 and 10B), which was not found in any other nematode pPolB2s (Extended Data Fig 1).

The pPolB2s of the marine nematode Polinton are highly diverged from the pPolB2s of the terrestrial nematode species. Among the known Polintons, the marine nematode pPolB2s showed the highest similarity to pPolB2s of mollusc Polintons as indicated by the phylogenetic tree topology (Fig 5B, Fig 6, and Extended Data Fig 8). Polintons of a mud snail (*B. attramentaria*) and a sea slug (*E. chlorotica*) encode the closest relatives of the marine nematode pPolB2 with the shared VSR endonuclease domain, an apparent synapomorphy of this subgroup of pPolB2 (Fig 6 and Extended Data Fig 8). The presence of the VSR domain along with the phylogenetic tree topology support an additional, independent inter-phylum horizontal gene transfer, in this case between Polintons of nematodes and molluscs Polintons that inhabit the same marine environments.

## Conclusions

In this work, we present the unexpected finding that nematode Polintons form two distinct groups that differ by encoding highly diverged pPolBs. The pPolB1 group is by far more widely spread and more abundant among the nematodes than the pPolB2 group suggesting that pPolB2 comparatively recently displaced pPolB1 in two distinct lineages of nematode Polintons at least one of which subsequently spread across nematode species. Phylogenetic analysis of a broad selection of protein-primed B family DNA polymerases suggested that pPolB2s originated from oomycete and mollusc Polintons independently of pPolB1. The DNA polymerase displacement then might have occurred by horizontal transfer of the pPolB2 gene, possibly, via infection of nematodes by oomycete and mollusc Polinton virus particles, with subsequent recombination with a resident nematode Polinton, resulting in the displacement of the DNA polymerase gene alone. The two independent incursions of pPolB2 subfamilies into the terrestrial and marine nematodes, from the oomycete and marine mollusc Polinton families respectively, indicate that the transfers are consistent with the native ecosystems of different groups of nematodes. In this context, we note that terrestrial nematodes can serve as hosts for natural oomycete infection, which could have facilitated the transfer^64,65^, whereas marine nematodes most likely shared environments, such as sediment, with prevalent marine molluscs. Although Polinton virions so far have not been observed, the conservation of the major and minor capsid proteins, the packaging ATPase and the maturation protease across the entire diversity of Polintons strongly suggests that such particles exist. Most likely, these virions mediate the Polinton spread, with the precise characteristics of genomic Polinton elements in a species group reflecting the historical environmental niche rather than the phylogeny. The findings presented in this paper are compatible with this prediction.

## Methods

### Search of Polintons through *Nematoda* genomes

Assembled genomes of *Nematoda* were downloaded from the NCBI genome assembly database by using the NCBI datasets (https://www.ncbi.nlm.nih.gov/datasets/) command-line tools. The taxon keyword “*Nematoda*” was used to download fasta format genome data.

Genome sequences were subjected to a tBLASTn search (with 0.001 as an *e*-value cutoff and 50 as a minimum bitscore). Query sequences (pPolB, INT, MCP, ATPase and Pro) for tBLASTn search were obtained from open reading frame (ORF) prediction by ORFfinder (https://www.ncbi.nlm.nih.gov/orffinder/) on *C. briggsae* Polinton-1 (Polinton-1_CB, WBTransposon00000832)^1^ that was downloaded from the WormBase (https://wormbase.org/). We filtered out candidate genetic blocks that had all five query proteins within 20 kb and gathered all of these segments with upstream and downstream buffers of 10 kb. Next, the candidates were further filtered out by the presence of terminal inverted repeats (TIR) that were detected using Inverted Repeat Finder (https://tandem-test.bu.edu/irf/irf.download.html)^66^. The 6 bases upstream and downstream of each candidate TIR were used to test whether a target site duplication (TSD) was present. Polinton candidate sequences were further validated by the presence of at least 5 Polinton proteins through ORF prediction on candidate sequences and protein-protein BLAST (BLASTp, bitscore cut-off: 100) of the ORFs against query Polinton proteins. A total of 266 Polintons from 66 nematode genomes (29 species) were identified and summarized in Extended Data Table 1.

### Multiple sequence alignment and phylogenetic tree analysis

Multiple sequence alignment for nematode pPolB proteins, *C. briggsae* Polinton proteins, and pPolB proteins found from each taxon group was performed using ClustalO^67,68^. Phylogenetic trees were generated with Maximum likelihood (ML) method by using IQ-TREE^69-71^, and mid-point rooted and visualized by ETE3 Python package^72^. Percent identity between protein sequences were calculated based on the pairwise comparison of sequences within the ClustalX environment^73^. Conserved motifs were searched from the multiple sequence alignment results and enrichment of amino acids were visualized by using LogoMaker Python package^74^.

For phylogenetic analyses of pPolBs from multiple taxa, pPolB protein sequences in each taxon group were clustered and a representative sequence was selected for each cluster by using CD-HIT^75^. Then, pPolBs were subjected to multiple sequence alignment by using ClustalO^67,68^. Gap regions in the alignment data were trimmed off with trimAl gappyout option^76^ and then phylogenetic trees were reconstructed by using ML method inferred by IQ-TREE v2.0.3 with ultrafast bootstrapping (1000 iterations)^69-71^ and with LG+F+R10 model as the best fitting model. The nodes of the phylogenetic tree were collapsed if all the pPolB leaves of the node were from same taxon groups. The phylogenetic trees were mid-point rooted and visualized by using ETE3 Python package^72^.

For the more global phylogenetic analysis, a dataset of pPolB sequences was retrieved from Krupovic and Koonin (2014)^10^. The dataset included pPolB sequences from bacterial tectiviruses, linear mitochondrial and cytoplasmic plasmids, adenoviruses and a limited number of bidnaviruses and Polintons. This dataset was supplemented with a greatly expanded set of pPolB sequences of Polintons, bidnaviruses and adintoviruses^25^. The dataset was then clustered to 50% identity over 80% of alignment length using MMseqs2^77,78^. The sequences were aligned using MAFFT with the G-INS-1 option^79^. Low-information content positions were removed using trimAl with the gappyout option^76^. The maximum likelihood phylogenetic tree was calculated using IQ-TREE v2.0.6^69^ with the best fitting substitution model determined for the data, which was LG+F+R10.

### Protein structure prediction and alignment

Structure prediction steps of *C. briggsae* pPolB1 or pPolB2 proteins were executed using MMSeqs2^77,78^ and AlphaFold2^56^ on ColabFold v1.5.2^57^ (https://colab.research.google.com/github/sokrypton/ColabFold/blob/main/AlphaFold2.ipynb). Pairwise structural alignment of predicted pPolB1 and pPolB2 structures was performed with FATCAT^80,81^ (https://www.rcsb.org/alignment) with default parameters. The structure was visualized using the PyMOL Molecular Graphics System, Version 2.5 Schrödinger, LLC.

### Polinton pPolB search through NCBI genome assemblies

We downloaded 7,897 genomes from 64 phyla of eukaryotes (except Viridiplantae or Chordata) available through the NCBI genome database as of late 2023 April (assembly source: genbank). To explore the distribution of nematode pPolB1 and pPolB2 homologs in the genomes, each fasta format genome file was indexed for tBLASTn search, and Polinton pPolB1 and pPolB2 protein sequences were searched through the genome using tBLASTn. To provide a stringent search, we filtered out small matches by setting a match length cutoff (650 amino acids for both pPolB1 and pPolB2). Then, the matched sequences with 20 kb both upstream and downstream were used to find potential ORFs for pPolB and other Polinton proteins. We further filtered candidates based on the length of predicted pPolB ORFs (a minimum ORF length: 900 amino acids). Subsequently, hmmsearch^82^ (http://hmmer.org/) was performed to detect homology between predicted ORFs and Polinton proteins. Profiles for the Polinton proteins (INT, MCP, and ATPase) were built from the multiple sequence alignments of each Polinton protein obtained from RepBase Polintons (https://www.girinst.org/repbase)^83,84^. To ensure that pPolB gene candidates are indeed Polinton polymerases, we selected pPolBs that co-detected with an INT protein nearby (see Extended Data Table 3 for pPolB-Polinton information). For each taxon group, copy numbers of pPolB1 and pPolB2 were normalized by the number of genomes that had at least a single copy of pPolB and visualized by using Python ETE3 package^72^.

## Acknowledgements

We thank Karen Lynn Artiles, Lamia Wahba, Massa Shoura, Matthew James McCoy, Orkan Ilbay, Jingxun Chen, Usman Enam, Emily Greenwald, Ivan Nikolay Zheludev, Drew Galls, and Janie Soo-hyun Kim for helpful discussion and their critical reading on the manuscript. This study was supported by the NIGMS grant (R35GM130366 to A.Z.F.), and by the Long-term Postdoctoral Fellowship from the Human Frontier Science Program (LT000329/2019-L) and the Bernard Cohen Postdoctoral Fellowship (to D-E.J.). S.S. is supported by the Stanford Cardiovascular Institute Summer Undergraduate Fellowship. E.V.K. is supported by the Intramural Research Program of the National Institutes of Health of the USA (National Library of Medicine).

## Supporting Figure legends

**Extended Data Fig 1.**
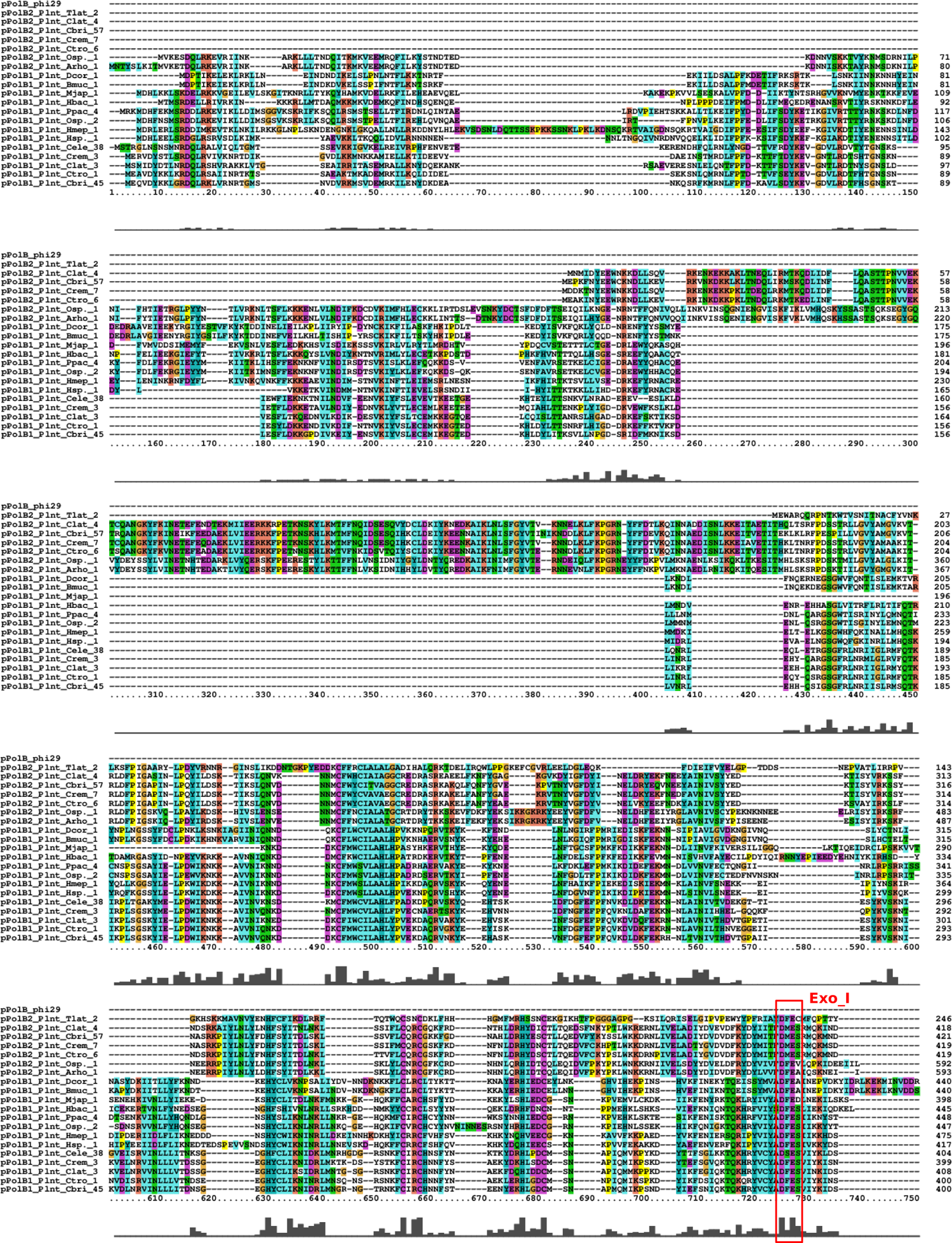
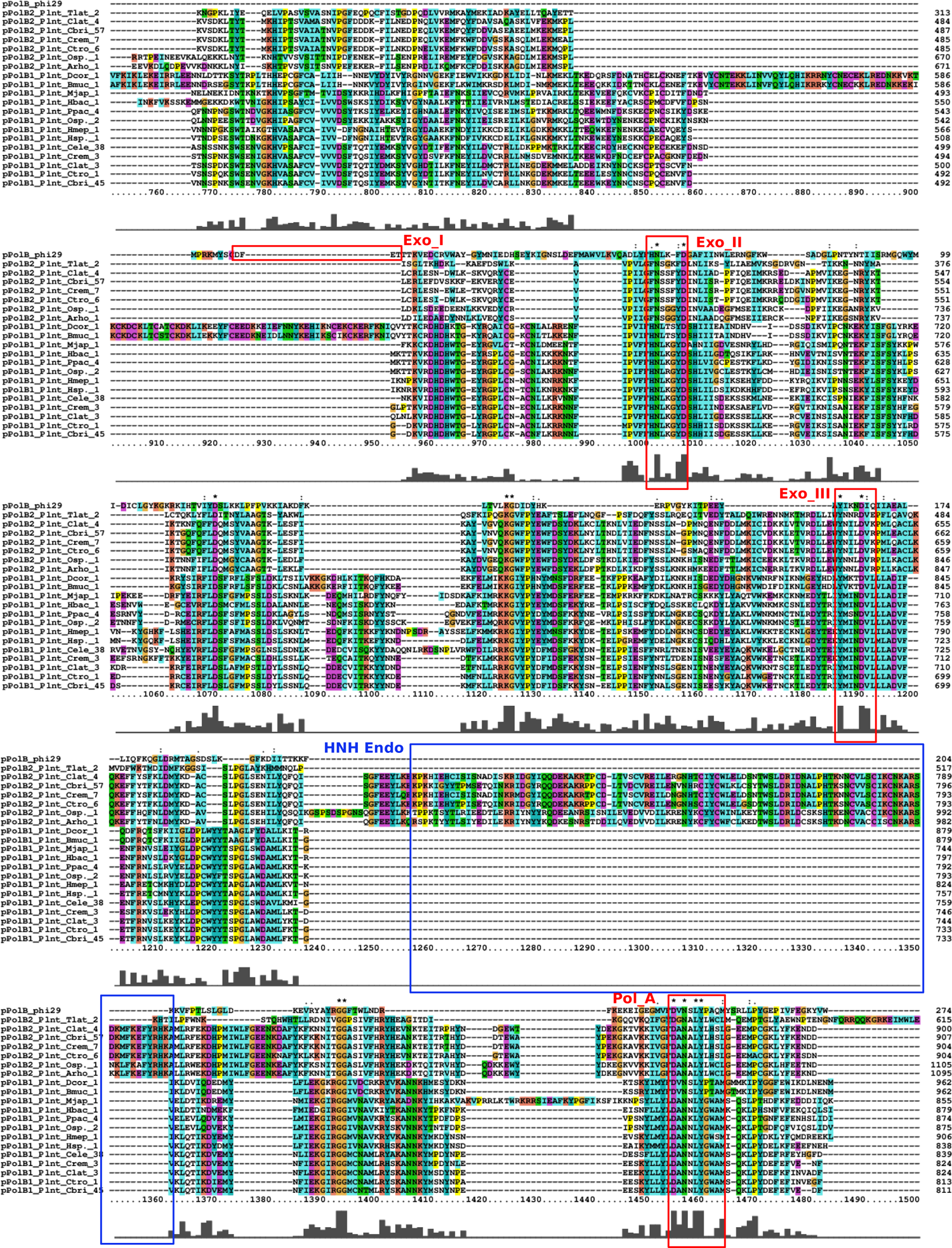
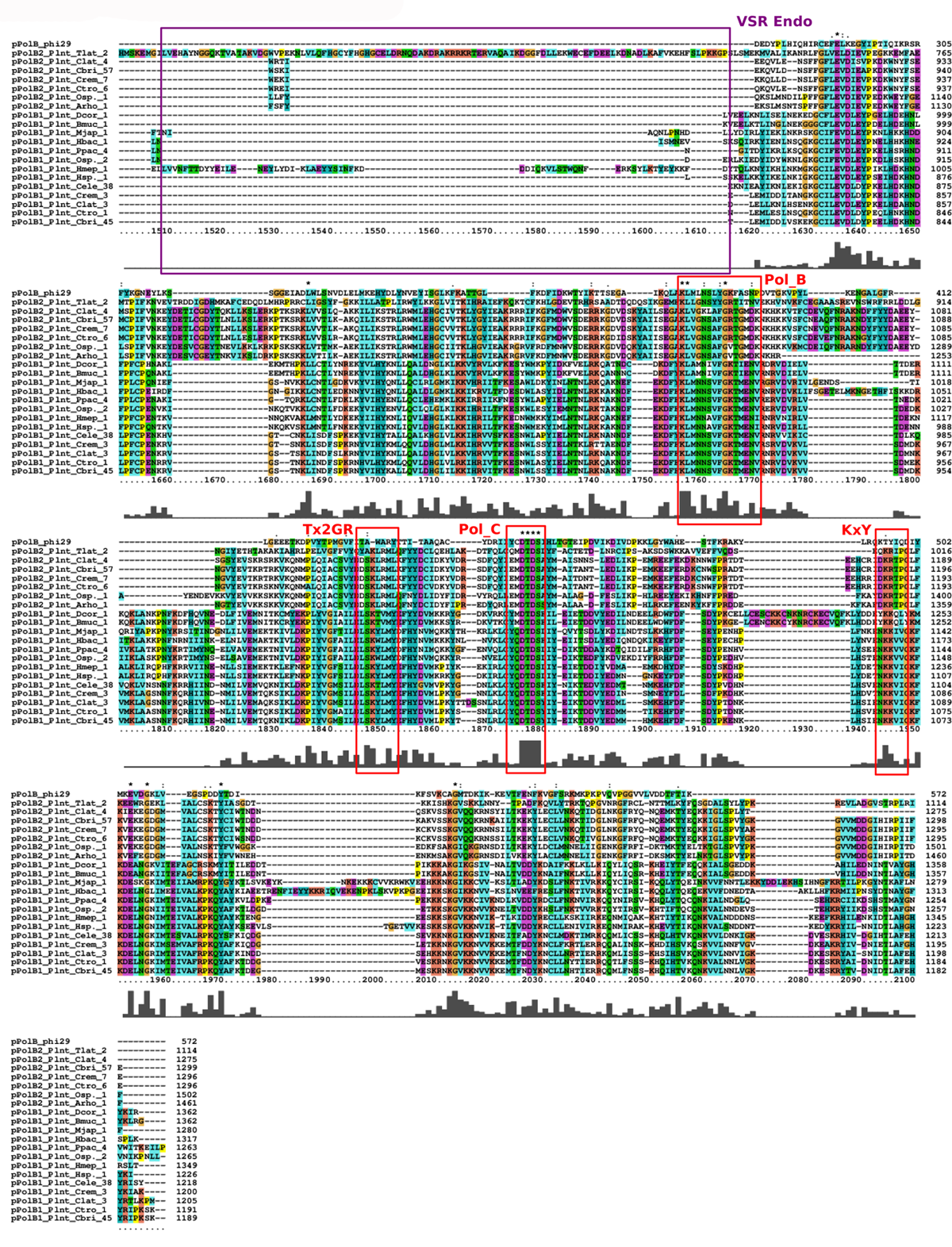
Nematode pPolB1 and pPolB2 have conserved exonuclease and polymerase motifs. Full-length multiple sequence alignment data of representative nematode pPolB proteins with catalytic motifs (Exo_I, Exo_II, Exo_III, Pol_A, Pol_B, Pol_C, KxY and Tx2GR) marked with red boxes. Nematode pPolB2 proteins contained HNH or VSR endonuclease domains marked with blue and purple boxes, respectively. Labels at the most left column show pPolB classes and Polinton unique identifiers (UI) (see Extended Data Table 1) that includes species information; Osp. for *Oscheius* species, Arho for *Auanema rhodensis*, Crem for *Caenorhabditis remanei*. Ctro for *Caenorhabditis tropicalis*, Cbri for *Caenorhabditis briggsae*, Clat for *Caenorhabditis latens*, Dcor for *Diploscapter coronatus*, Bmuc for *Bursaphelenchus mucronatus*, Cele for *Caenorhabditis elegans*, Hmep for *Halicephalobus mephisto*, Hsp. for *Halicephalobus* species, Ppac for *Pristionchus pacificus*, Hbac for *Heterorhabditis bacteriophora*, Mjap for *Micoletzkya japonica*, and Tlat for *Trissonchulus latispiculum*.

**Extended Data Fig 2.**
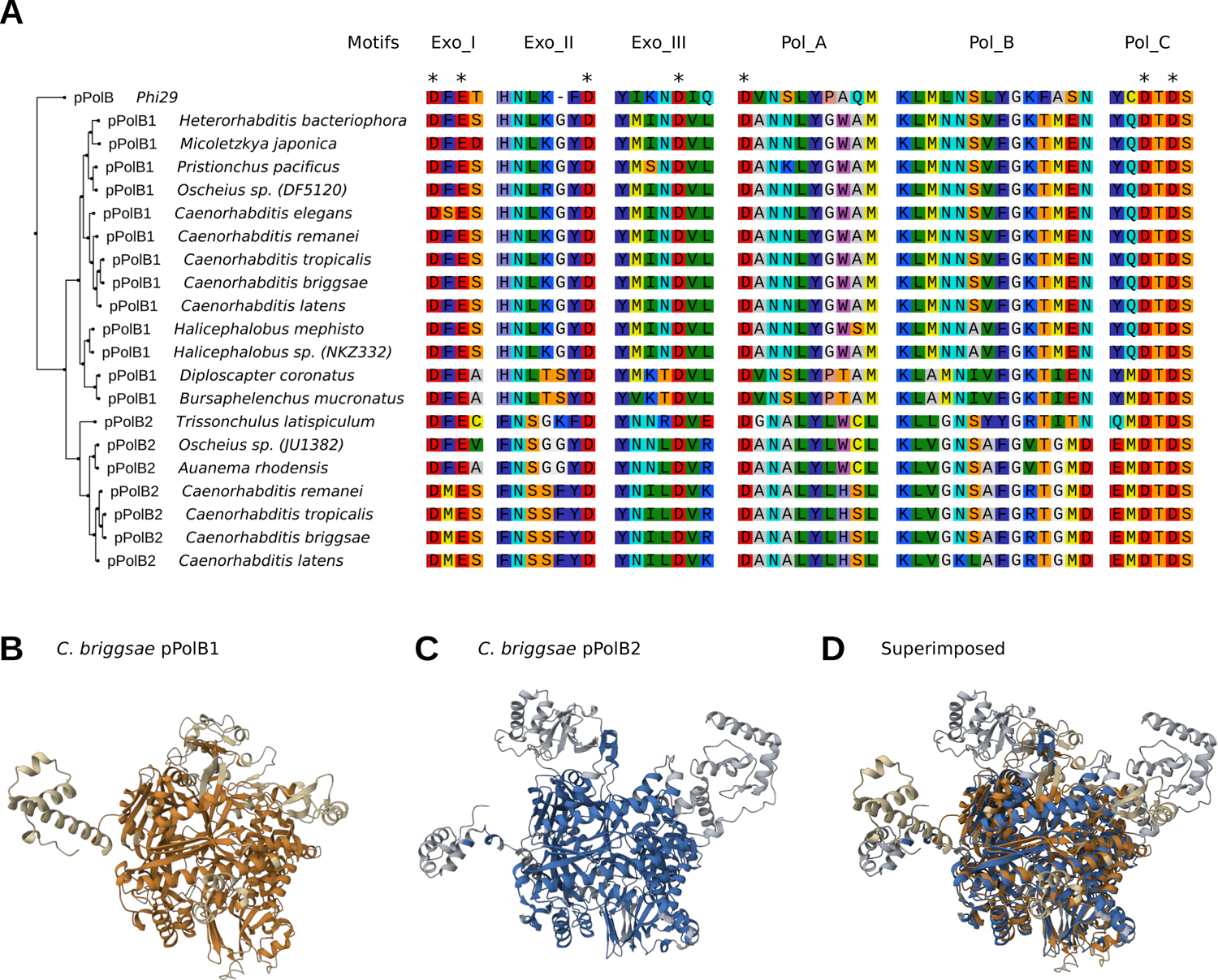
pPolB1 and pPolB2 proteins have closely similar structures with conserved catalytic motifs. **A.** Multiple sequence alignment (right panels) of pPolB1 and pPolB2 proteins from 19 nematode species. The phylogenetic tree (left panels) is equivalent to that in Fig 1A with the minor addition of the outgroup Phi29 phage DNA polymerase). Six conserved pPolB protein motifs (Exo_I, Exo_II, Pol_A, Pol_B, and Pol_C) are selectively shown (See Extended Data Fig 1 for a full alignment result). **B-D**. show AlphaFold2 predictions of *C. briggsae* pPolB1 (orange, **B**), pPolB2 (blue, **C**) and a superimposed structure (RMSD: 4.22, TM-score: 0.53) (**D**) of pPolB1 and pPolB2 constructed by structural alignment. Dark and light colors indicate aligned and non-aligned positions within these putative structures, respectively.

**Extended Data Fig 3.**
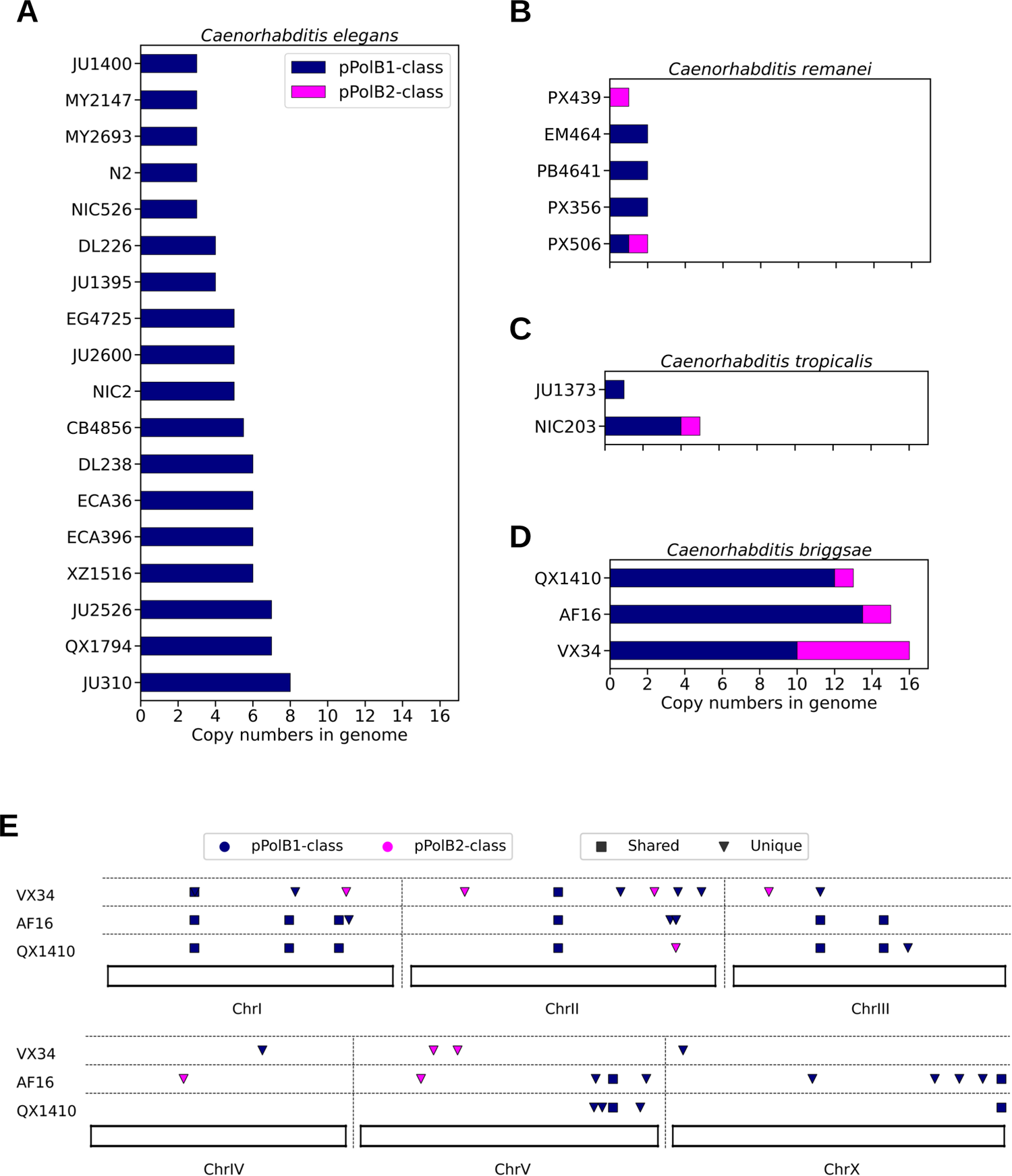
intra-species copy number and position variations of *Caenorhabditis* Polintons. **A-D.** Bar graphs showing average copy numbers per genome of pPolB1-class (navy) and pPolB2-class (magenta) full-length Polintons in wild-type strains of *C. elegans* (A), *C. remanei* (**B**), *C. tropicalis* (**C**) and *C. briggsae* (**D**). **E.** Insertion positions of pPolB1-(navy) and pPolB2-class (magenta) Polintons from three *C. briggsae* strains (VX34, AF16, and QX1410) at the chromosome scale. Dot shapes indicate whether the positions were found as unique (inversed triangle) or shared (square) in at least two strains.

**Extended Data Fig 4.**
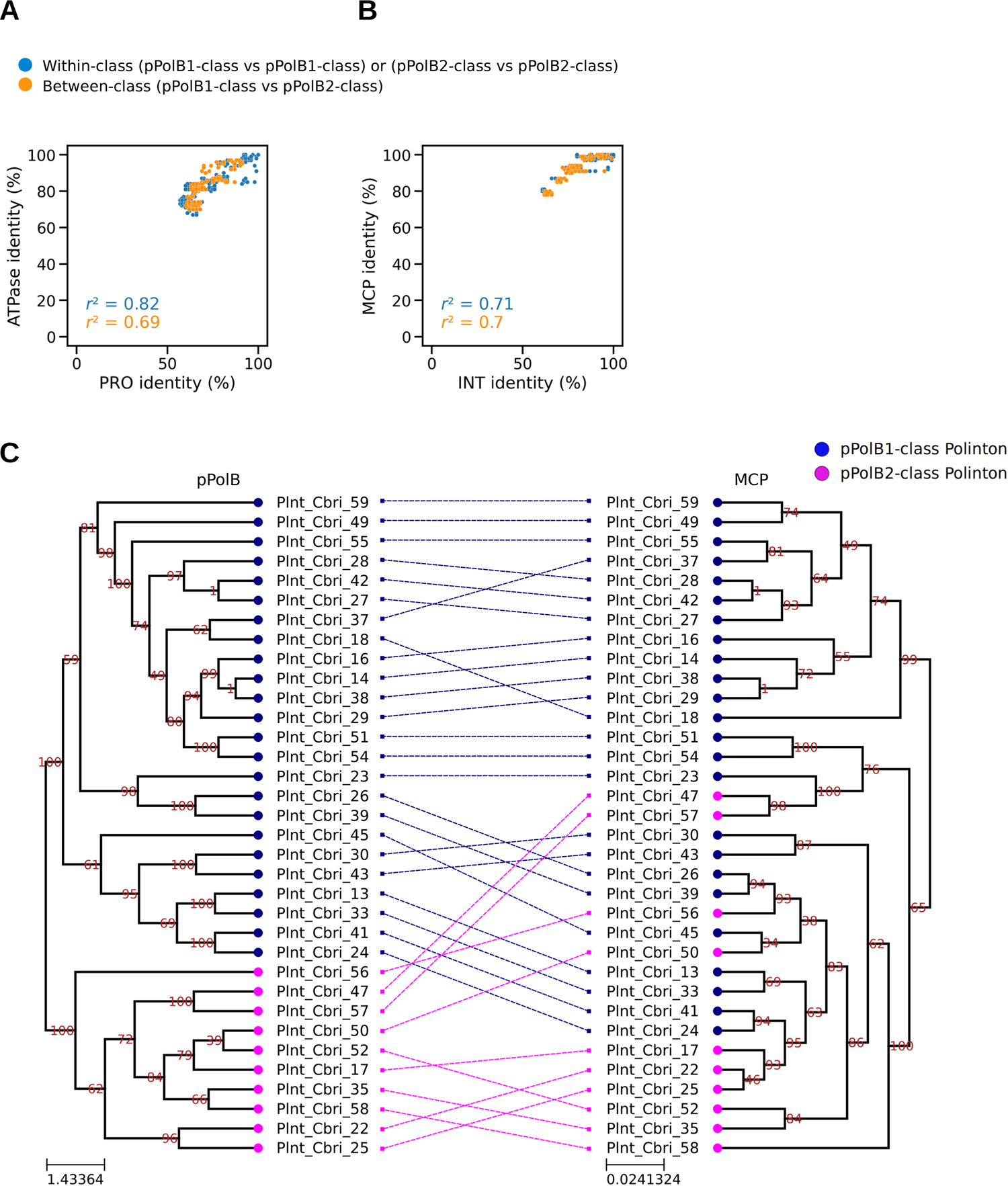
Polinton genes diverge together. **A-B**. Scatter plots showing percent (%) amino acid identity of a protein at each axis from *C. briggsae* Polinton species; ATPase and PRO (**A**), INT and MCP (**B**). Each dot indicates a pair of two *C. briggsae* Polintons, whose proteins indicated at the X and Y axes were aligned, respectively, for calculating % identity. Dot colors indicate whether two Polintons in a pair for comparison are within the same groups of Polintons (within pPolB-class, blue) or from different groups (between pPolB-class, orange). **C.** A tanglegram showing the relationship between two phylogenetic trees constructed from multiple sequence alignment of *C. briggsae* pPolB1/2 (left panel) and MCP (right panel) proteins. The colors of the nodes indicate pPolB1-class (navy) and pPolB2-class (magenta). Dashed lines between two trees denote the correspondence of pPolB and MCP proteins found from the same Polinton.

**Extended Data Fig 5.**
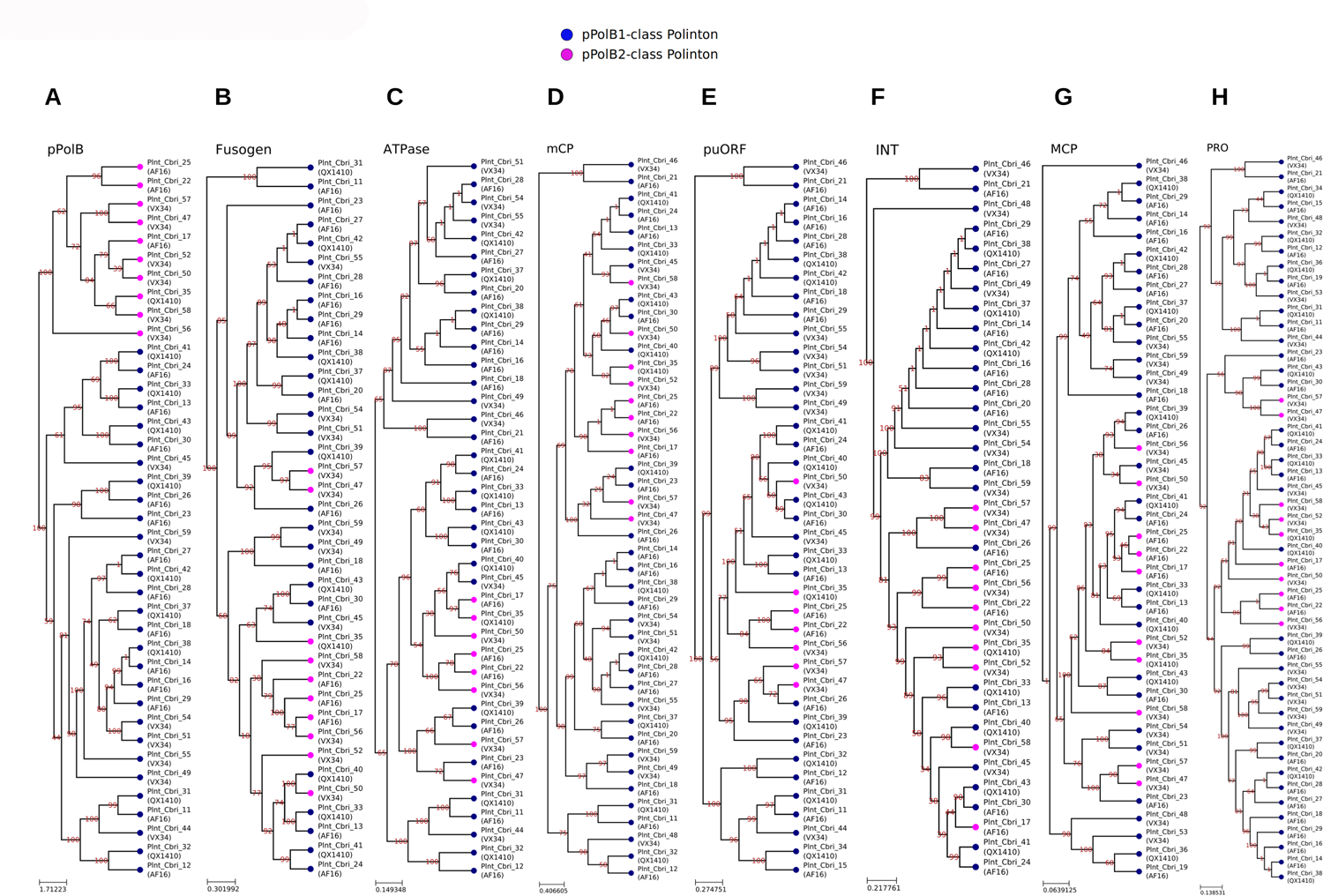
Phylogenetic analysis suggests acquisition of two classes of pPolB in *C. briggsae* Polintons during evolution. **A-H**. Phylogenetic trees were constructed by maximum likelihood method on multiple sequence alignment of pPolB (**A**), Fusogen (**B**), ATPase (**C**), mCP (**D**), puORF (**E**), INT (**F**), MCP (**G**) and PRO (**H**) proteins found in *C. briggsae* Polintons. Navy and magenta colors of tree nodes indicates pPolB1- and pPolB2-class Polintons, respectively, where the proteins are encoded. The numbers on each branch indicates bootstrap supporting values.

**Extended Data Fig 6.**
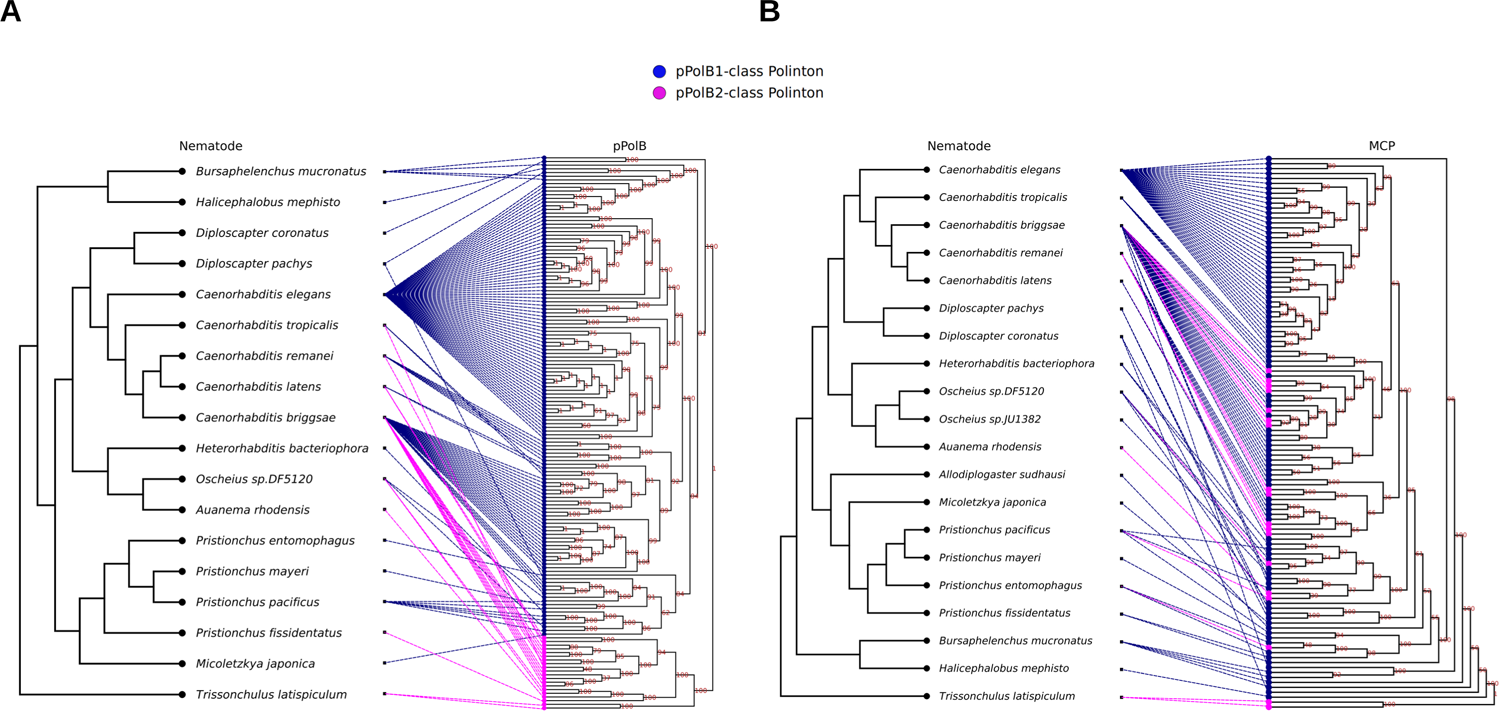
Evidence for horizontal gene transfer mediated by Polintons across nematode species. **A-B.** Tanglegrams that show the link between phylogenetic trees of nematode species (left) and Polinton pPolB (**A**) and MCP (**B**) proteins. The colors of tree nodes indicate pPolB1-class (navy) and pPolB2-class (magenta). Dashed lines between two trees denote the correspondence of pPolB (**A**) or MCP (**B**) proteins found from the nematode species (navy and magenta for the links from species to pPolB1- and pPolB2-class Polintons, respectively). Subsets of closely related PolBs and/or MCPs were found across distant nematode species, suggesting either that these Polintons originate from a common Polinton ancestor or that horizontal gene transfer event occurred across different nematode species.

**Extended Data Fig 7.**
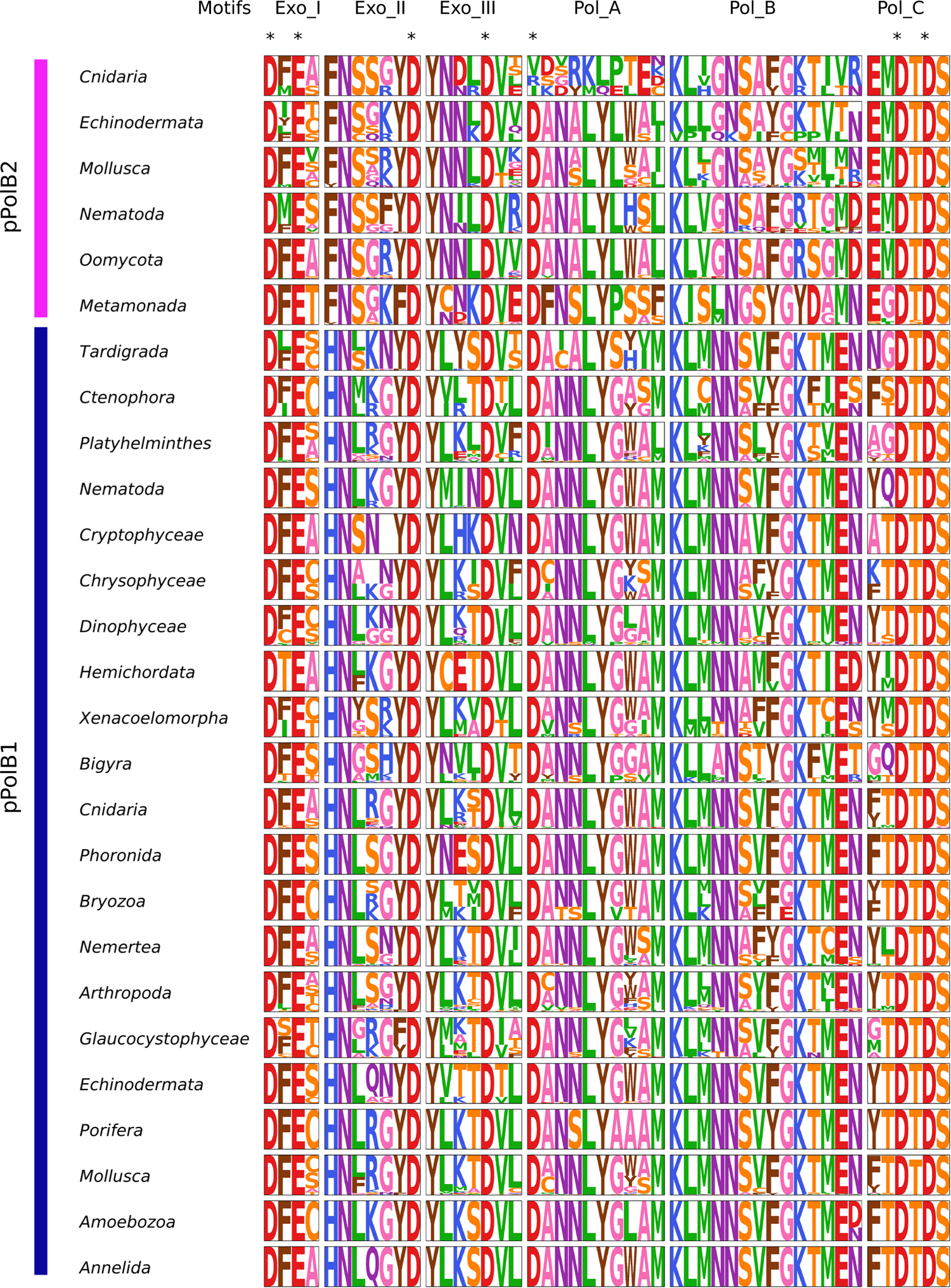
Polinton pPolBs have conserved exonuclease and polymerase domains. Conserved exonuclease (Exo_I, Exo_II, and Exo_III) and polymerase (Pol_A, Pol_B, and Pol_C) motifs for pPolBs found from each taxon groups were searched through multiple sequence alignments and are shown with graphical representation of the conserved amino acids.

**Extended Data Fig 8.**
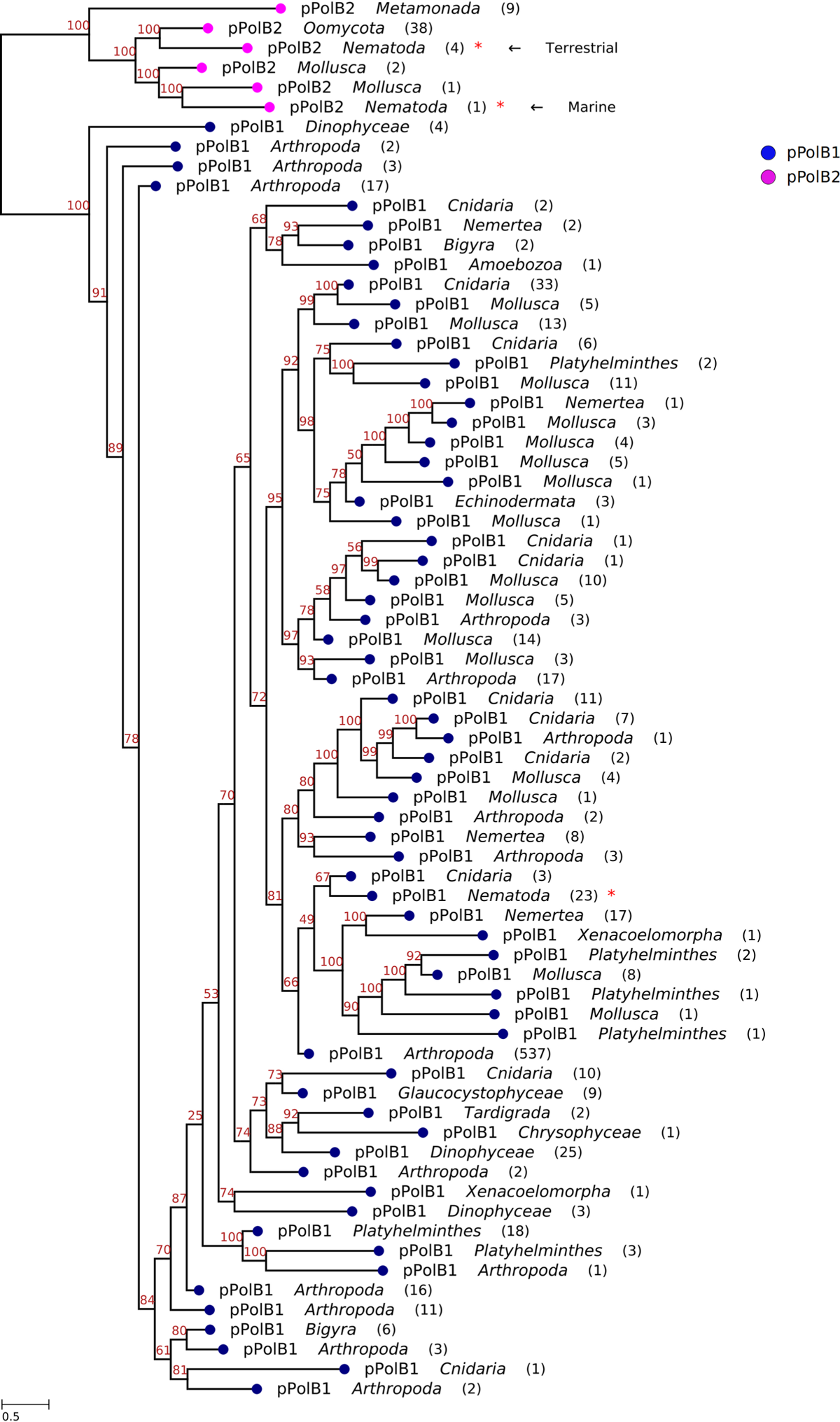
Monophyletic groups of Polinton pPolB1 and pPolB2 families. A phylogenetic tree of representative pPolB1 and pPolB2 proteins from multiple taxon groups was built by using IQ-TREE-inferred maximum likelihood method on multiple sequence alignment of the proteins (see Method for details). Bootstrap supporting values for each branch were shown on the top position of the branches. Navy and magenta colors indicate pPolB1 and pPolB2, respectively. The numbers in parentheses denote the number of pPolB proteins in each node of taxon group used for the multiple sequence alignment. Nematode pPolB2 groups found from the terrestrial and marine species are marked with red arrows.

**Extended Data Fig 9.**
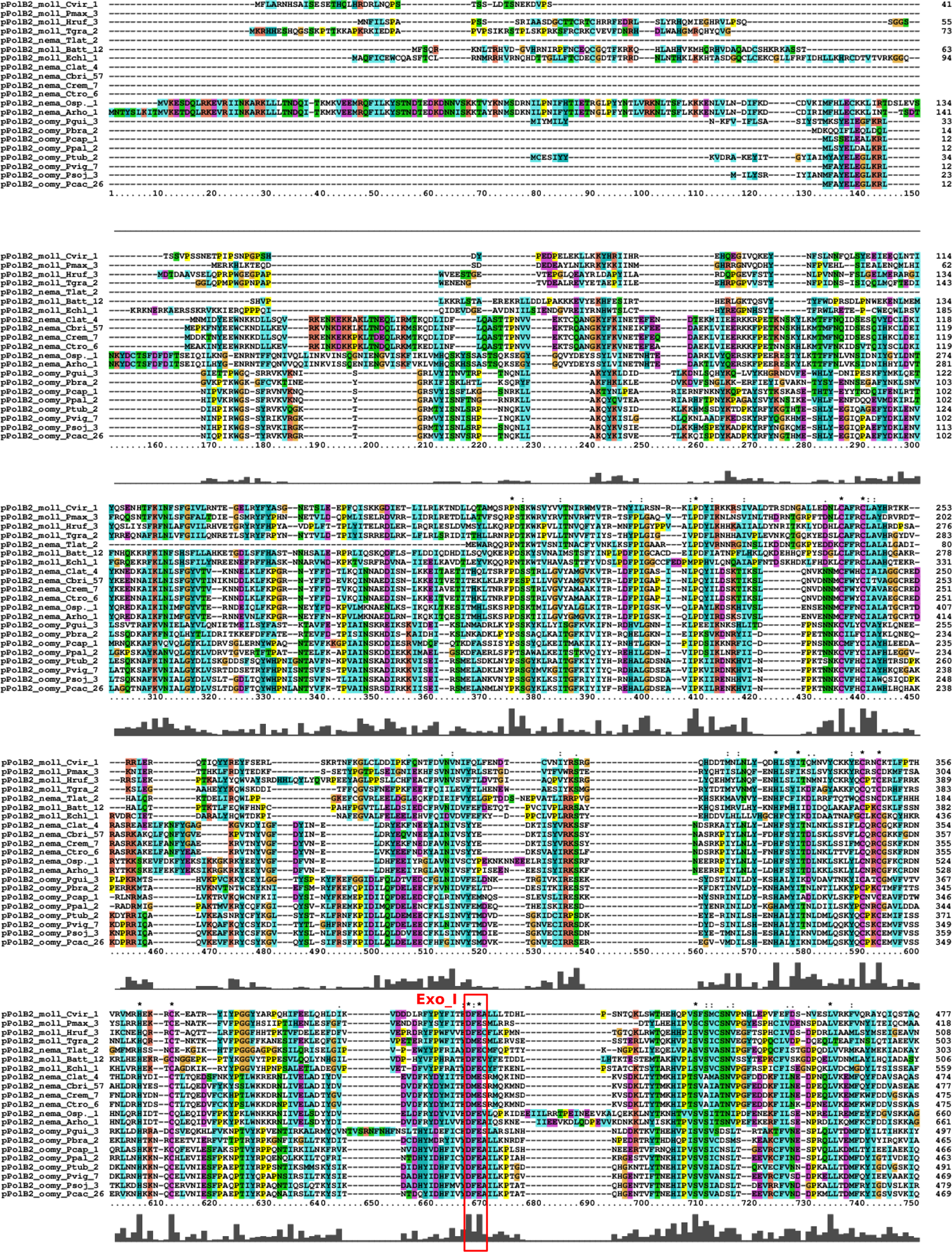
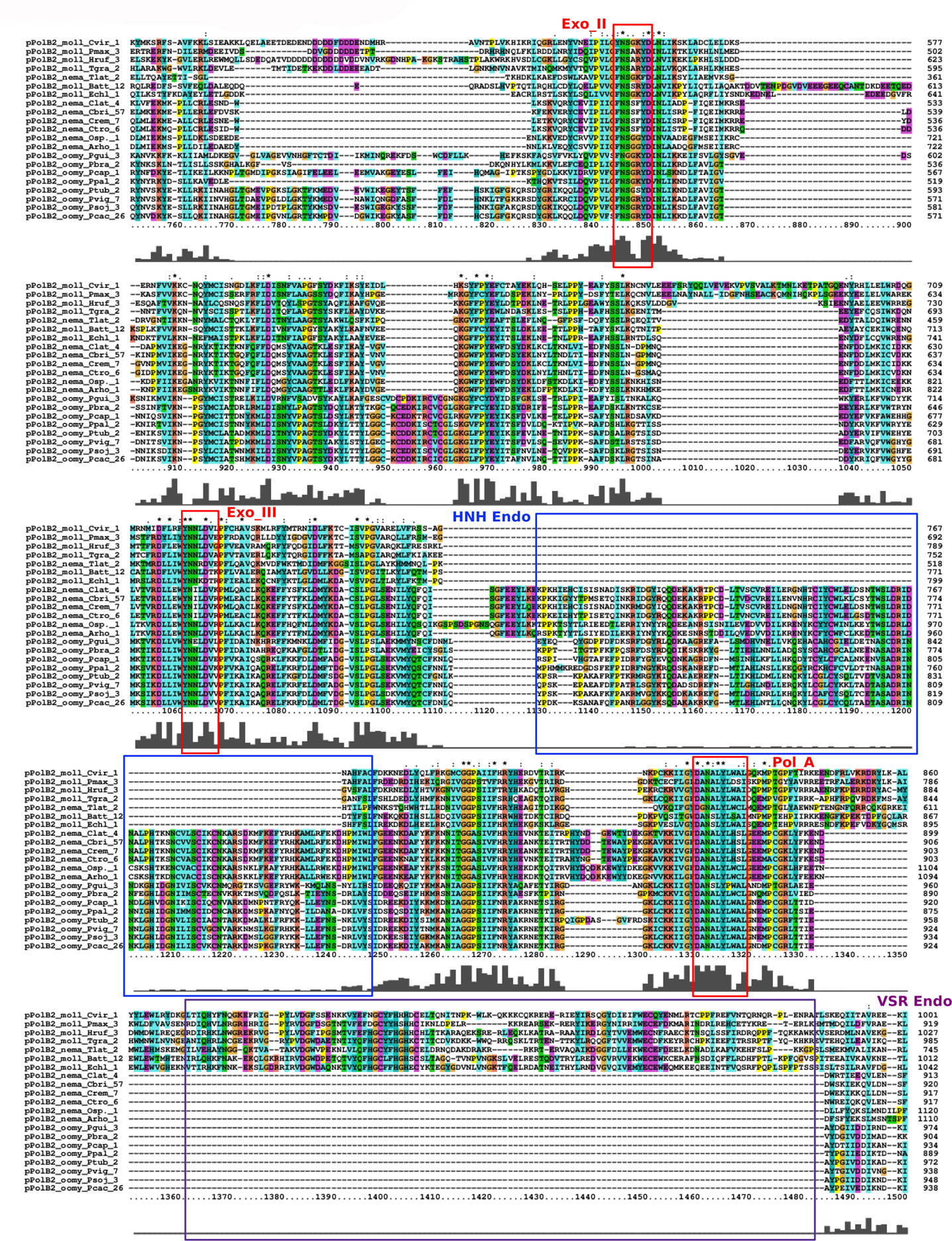
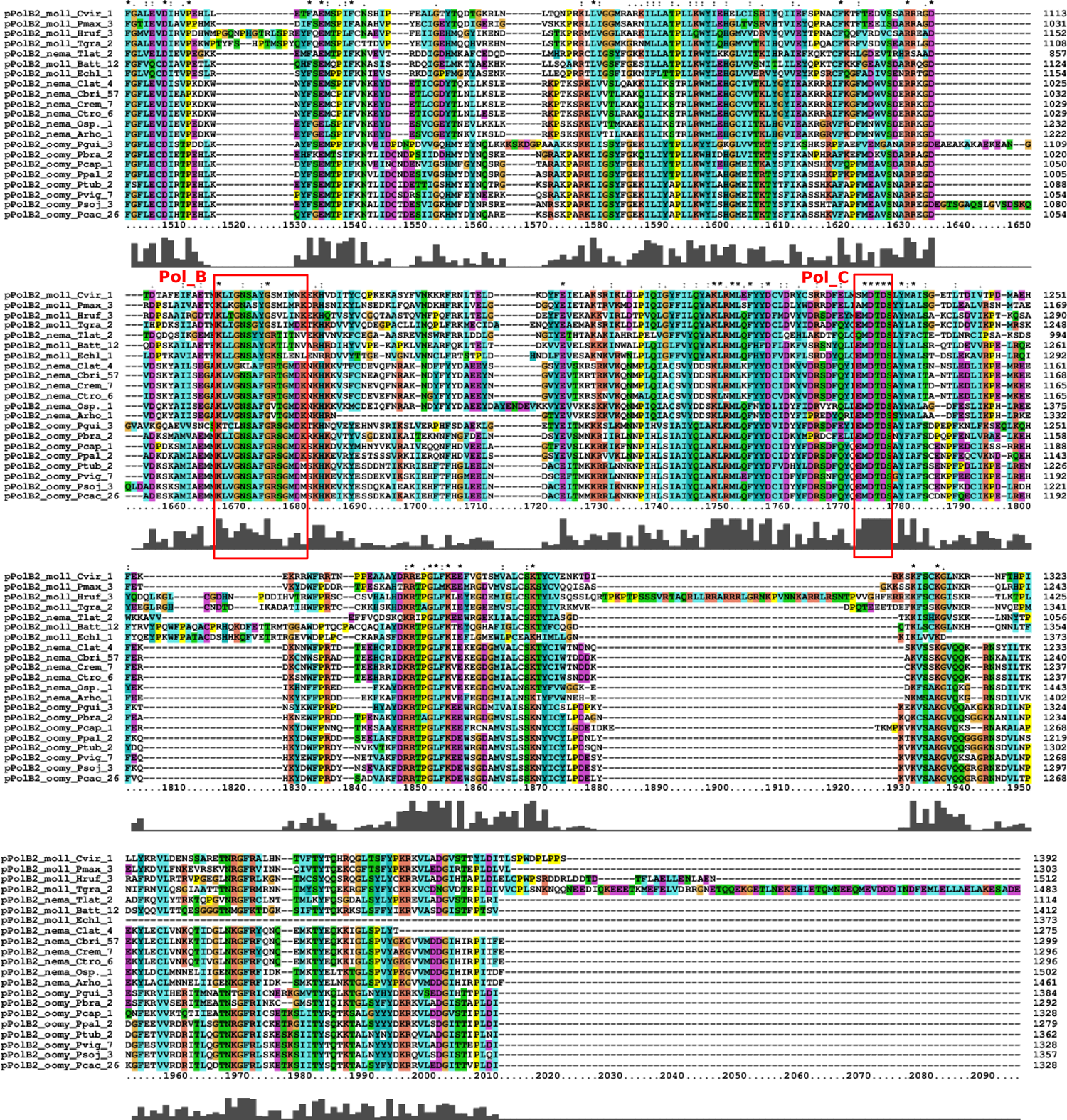
Conserved Polinton pPolB domains from three distinct phyla. Full-length multiple sequence alignment data of representative pPolB2 proteins from three distinct phyla; *Nematoda*, *Oomycota*, and *Mollusca*. Red boxes indicate conserved catalytic motifs for exonuclease (Exo_I, Exo_II, Exo_III) and polymerase (Pol_A, Pol_B, Pol_C) activity. The conserved HNH endonuclease domain marked with a blue box is found in terrestrial nematode and oomycete pPolB2s but not in mollusc pPolB2s. In contrast, The VSR endonuclease domain marked with a purple box was conserved in a marine nematode pPolB2 (nema_Tlat_2) and mollusc pPolB2 proteins. Labels at the most left column show pPolB2 unique identifier that includes phylum and species information; moll for *Mollusca*, nema fo*r Nematoda*, oomy for *Oomycota*, Cvir for *Crassorstrea virginica*, Batt for *Batillaria attramentaria*, Hruf for *Haliotis rufescens*, Echl for *Elysia chlorotica*, Pmax for *Pecten maximus*, Tgra for *Tegillarca granosa*, Tlat for *Trissonchulus latispiculum*, Osp. for *Oscheius* species, Arho for *Auanema rhodensis*, Crem for *Caenorhabditis remanei*, Clat for *Caenorhabditis latens*, Ctro for *Caenorhabditis tropicalis*, Cbri for *Caenorhabditis briggsae*, Aeut for Aphanomyces euteiches, Lgig for Lottia gigantea, Pcac for *Phytophthora cactorum*, Pcap for *Phytophthora capsici*, Pfra for *Phytophthora fragariae*, Plat for *Phytophthora lateralis*, Ppal for *Phytophthora palmivora*, Psoj for *Phytophthora sojae*, Ptub for *Phytophthora tubulina*, Pvig for *Phytophthora vignae*, Pbra for *Pythium brassicum*, Pgui for *Pythium guiyangense*.

**Extended Data Fig 10.**
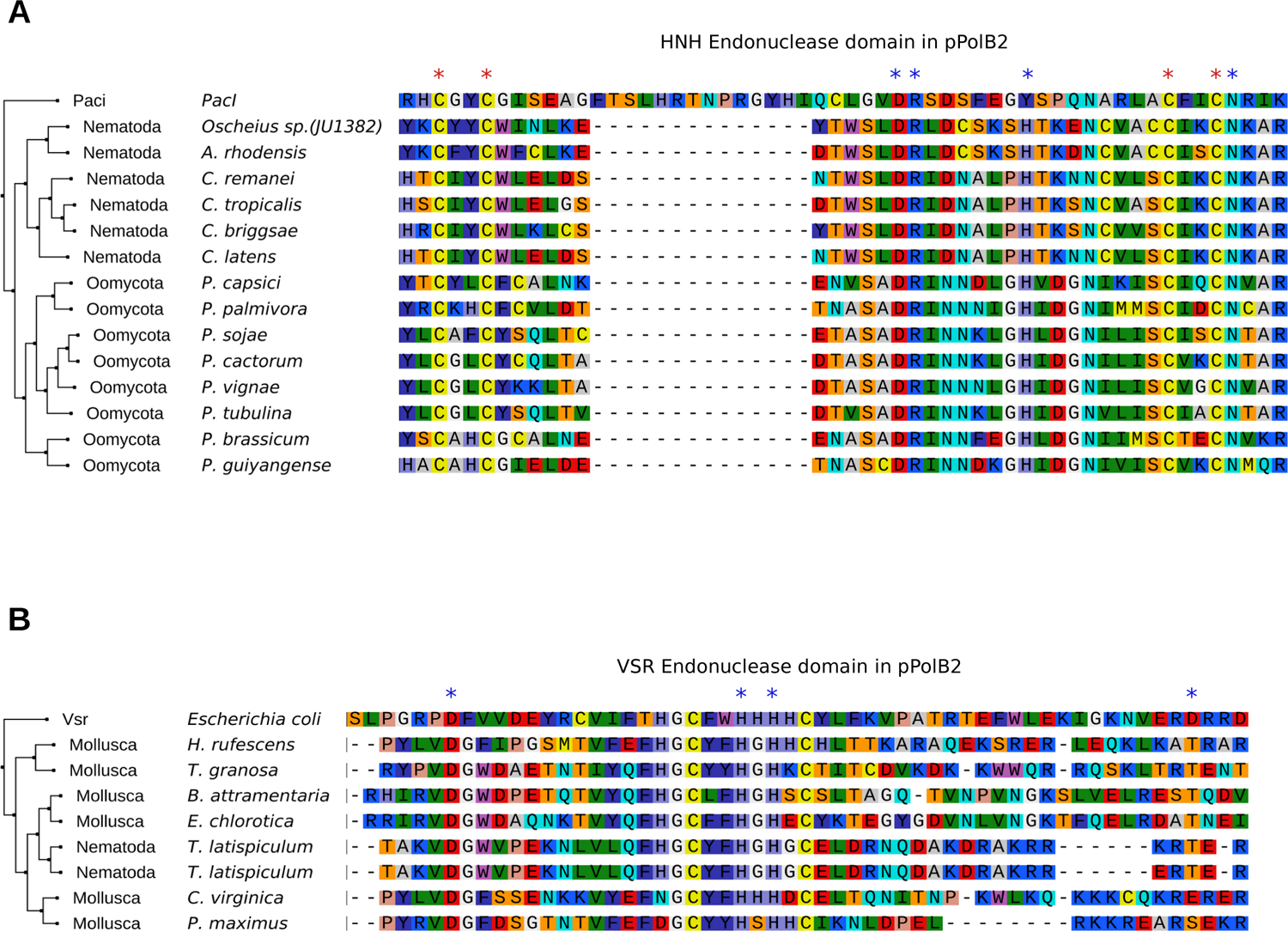
HNH and VSR domains of pPolB2s appear functional. **A.** A phylogenetic tree (left) and multiple sequence alignment (right) of HNH endonuclease domains found in Polinton pPolB2 proteins of nematode and oomycete species with PacI bacterial HNH endonuclease found in *Pseudomonas alcaligenes*. Red and blue asterisks indicate zinc-binding Cys residues and catalytic residues for PacI^59^, both of which are conserved across nematode and oomycete pPolB2 proteins except a Tyr residue. **B.** A phylogenetic tree (left) and multiple sequence alignment (right) of VSR endonuclease domains found in Polinton pPolB2 proteins of marine nematode and mollusc species with *E. coli* VSR protein. Blue asterisks indicate catalytically important residues for *E. coli* VSR endonuclease activity^87^.

## References

1 Kapitonov, V. V. & Jurka, J. Self-synthesizing DNA transposons in eukaryotes. Proc Natl Acad Sci U S A 103, 4540–4545, doi:10.1073/pnas.0600833103 (2006).

2 Pritham, E. J., Putliwala, T. & Feschotte, C. Mavericks, a novel class of giant transposable elements widespread in eukaryotes and related to DNA viruses. Gene 390, 3–17, doi:10.1016/j.gene.2006.08.008 (2007).

3 Feschotte, C. & Pritham, E. J. DNA transposons and the evolution of eukaryotic genomes. Annu Rev Genet 41, 331–368, doi:10.1146/annurev.genet.40.110405.090448 (2007).

4 Jurka, J., Kapitonov, V. V., Kohany, O. & Jurka, M. V. Repetitive sequences in complex genomes: structure and evolution. Annu Rev Genomics Hum Genet 8, 241–259, doi:10.1146/annurev.genom.8.080706.092416 (2007).

5 Krupovic, M. & Koonin, E. V. Polintons: a hotbed of eukaryotic virus, transposon and plasmid evolution. Nat Rev Microbiol 13, 105–115, doi:10.1038/nrmicro3389 (2015).

6 Feschotte, C. & Pritham, E. J. Non-mammalian c-integrases are encoded by giant transposable elements. Trends Genet 21, 551–552, doi:10.1016/j.tig.2005.07.007 (2005).

7 Krupovic, M., Bamford, D. H. & Koonin, E. V. Conservation of major and minor jelly-roll capsid proteins in Polinton (Maverick) transposons suggests that they are bona fide viruses. Biol Direct 9, 6, doi:10.1186/1745-6150-9-6 (2014).

8 Gao, X. & Voytas, D. F. A eukaryotic gene family related to retroelement integrases. Trends Genet 21, 133–137, doi:10.1016/j.tig.2005.01.006 (2005).

9 Krupovic, M. & Koonin, E. V. Self-synthesizing transposons: unexpected key players in the evolution of viruses and defense systems. Curr Opin Microbiol 31, 25–33, doi:10.1016/j.mib.2016.01.006 (2016).

10 Krupovic, M. & Koonin, E. V. Evolution of eukaryotic single-stranded DNA viruses of the Bidnaviridae family from genes of four other groups of widely different viruses. Sci Rep 4, 5347, doi:10.1038/srep05347 (2014).

11 Koonin, E. V., Krupovic, M. & Yutin, N. Evolution of double-stranded DNA viruses of eukaryotes: from bacteriophages to transposons to giant viruses. Ann N Y Acad Sci 1341, 10–24, doi:10.1111/nyas.12728 (2015).

12 Haapa-Paananen, S., Wahlberg, N. & Savilahti, H. Phylogenetic analysis of Maverick/Polinton giant transposons across organisms. Mol Phylogenet Evol 78, 271–274, doi:10.1016/j.ympev.2014.05.024 (2014).

13 Novick, P. A., Smith, J. D., Floumanhaft, M., Ray, D. A. & Boissinot, S. The evolution and diversity of DNA transposons in the genome of the Lizard Anolis carolinensis. Genome Biol Evol 3, 1–14, doi:10.1093/gbe/evq080 (2011).

14 da Silva, A. F., Dezordi, F. Z., Loreto, E. L. S. & Wallau, G. L. Drosophila parasitoid wasps bears a distinct DNA transposon profile. Mob DNA 9, 23, doi:10.1186/s13100-018-0127-2 (2018).

15 Shao, F., Han, M. & Peng, Z. Evolution and diversity of transposable elements in fish genomes. Sci Rep 9, 15399, doi:10.1038/s41598-019-51888-1 (2019).

16 Klai, K. et al. Screening of Helicoverpa armigera Mobilome Revealed Transposable Element Insertions in Insecticide Resistance Genes. Insects 11, doi:10.3390/insects11120879 (2020).

17 Bellas, C. et al. Large-scale invasion of unicellular eukaryotic genomes by integrating DNA viruses. Proc Natl Acad Sci U S A 120, e2300465120, doi:10.1073/pnas.2300465120 (2023).

18 Chase, E. E., Desnues, C. & Blanc, G. Integrated viral elements suggest the dual lifestyle of Tetraselmis spp. Polinton-like viruses. Virus Evol 8, veac068, doi:10.1093/ve/veac068 (2022).

19 Fischer, M. G. & Suttle, C. A. A virophage at the origin of large DNA transposons. Science 332, 231–234, doi:10.1126/science.1199412 (2011).

20 Fischer, M. G. & Hackl, T. Host genome integration and giant virus-induced reactivation of the virophage mavirus. Nature 540, 288–291, doi:10.1038/nature20593 (2016).

21 Roux, S. et al. Updated Virophage Taxonomy and Distinction from Polinton-like Viruses. Biomolecules 13, doi:10.3390/biom13020204 (2023).

22 Yutin, N., Shevchenko, S., Kapitonov, V., Krupovic, M. & Koonin, E. V. A novel group of diverse Polinton-like viruses discovered by metagenome analysis. BMC Biol 13, 95, doi:10.1186/s12915-015-0207-4 (2015).

23 Bellas, C. M. & Sommaruga, R. Polinton-like viruses are abundant in aquatic ecosystems. Microbiome 9, 13, doi:10.1186/s40168-020-00956-0 (2021).

24 Roitman, S. et al. Isolation and infection cycle of a polinton-like virus virophage in an abundant marine alga. Nat Microbiol 8, 332–346, doi:10.1038/s41564-022-01305-7 (2023).

25 Starrett, G. J. et al. Adintoviruses: a proposed animal-tropic family of midsize eukaryotic linear dsDNA (MELD) viruses. Virus Evol 7, veaa055, doi:10.1093/ve/veaa055 (2021).

26 Bessereau, J. L. Transposons in C. elegans. WormBook, 1–13, doi:10.1895/wormbook.1.70.1 (2006).

27 Laricchia, K. M., Zdraljevic, S., Cook, D. E. & Andersen, E. C. Natural Variation in the Distribution and Abundance of Transposable Elements Across the Caenorhabditis elegans Species. Mol Biol Evol 34, 2187–2202, doi:10.1093/molbev/msx155 (2017).

28 De Ley, P. A quick tour of nematode diversity and the backbone of nematode phylogeny. WormBook, 1–8, doi:10.1895/wormbook.1.41.1 (2006).

29 Vanreusel, A., De Groote, A., Gollner, S. & Bright, M. Ecology and biogeography of free-living nematodes associated with chemosynthetic environments in the deep sea: a review. PLoS One 5, e12449, doi:10.1371/journal.pone.0012449 (2010).

30 Nigon, V. M. & Felix, M. A. History of research on C. elegans and other free-living nematodes as model organisms. WormBook 2017, 1–84, doi:10.1895/wormbook.1.181.1 (2017).

31 Frezal, L. & Felix, M. A. C. elegans outside the Petri dish. Elife 4, doi:10.7554/eLife.05849 (2015).

32 Hodda, M. Phylum Nematoda: a classification, catalogue and index of valid genera, with a census of valid species. Zootaxa 5114, 1–289, doi:10.11646/zootaxa.5114.1.1 (2022).

33 Majdi, N. & Traunspurger, W. Free-living nematodes in the freshwater food web: a review. J Nematol 47, 28–44 (2015).

34 Schulenburg, H. & Felix, M. A. The Natural Biotic Environment of Caenorhabditis elegans. Genetics 206, 55–86, doi:10.1534/genetics.116.195511 (2017).

35 Felix, M. A. et al. Natural and experimental infection of Caenorhabditis nematodes by novel viruses related to nodaviruses. PLoS Biol 9, e1000586, doi:10.1371/journal.pbio.1000586 (2011).

36 Zhang, F. et al. Caenorhabditis elegans as a Model for Microbiome Research. Front Microbiol 8, 485, doi:10.3389/fmicb.2017.00485 (2017).

37 Felix, M. A. & Wang, D. Natural Viruses of Caenorhabditis Nematodes. Annu Rev Genet 53, 313–326, doi:10.1146/annurev-genet-112618-043756 (2019).

38 Pulavarty, A., Egan, A., Karpinska, A., Horgan, K. & Kakouli-Duarte, T. Plant Parasitic Nematodes: A Review on Their Behaviour, Host Interaction, Management Approaches and Their Occurrence in Two Sites in the Republic of Ireland. Plants (Basel) 10, doi:10.3390/plants10112352 (2021).

39 Bekal, S., Domier, L. L., Niblack, T. L. & Lambert, K. N. Discovery and initial analysis of novel viral genomes in the soybean cyst nematode. J Gen Virol 92, 1870–1879, doi:10.1099/vir.0.030585-0 (2011).

40 Bekal, S. et al. A novel flavivirus in the soybean cyst nematode. J Gen Virol 95, 1272–1280, doi:10.1099/vir.0.060889-0 (2014).

41 Lin, J. et al. A novel picornavirus-like genome from transcriptome sequencing of sugar beet cyst nematode represents a new putative genus. J Gen Virol 99, 1418–1424, doi:10.1099/jgv.0.001139 (2018).

42 Ruark, C. L. et al. Soybean cyst nematode culture collections and field populations from North Carolina and Missouri reveal high incidences of infection by viruses. PLoS One 12, e0171514, doi:10.1371/journal.pone.0171514 (2017).

43 Vieira, P. & Nemchinov, L. G. A novel species of RNA virus associated with root lesion nematode Pratylenchus penetrans. J Gen Virol 100, 704–708, doi:10.1099/jgv.0.001246 (2019).

44 Franz, C. J., Zhao, G., Felix, M. A. & Wang, D. Complete genome sequence of Le Blanc virus, a third Caenorhabditis nematode-infecting virus. J Virol 86, 11940, doi:10.1128/JVI.02025-12 (2012).

45 Williams, S. H. et al. Discovery of two highly divergent negative-sense RNA viruses associated with the parasitic nematode, Capillaria hepatica, in wild Mus musculus from New York City. J Gen Virol 100, 1350–1362, doi:10.1099/jgv.0.001315 (2019).

46 Brown, D. J. F., Robertson, W. M. & Trudgill, D. L. Transmission of Viruses by Plant Nematodes. Annual Review of Phytopathology 33, 223–249, doi:DOI 10.1146/annurev.py.33.090195.001255 (1995).

47 Hess, R. T. & Poinar, G. O., Jr. Iridoviruses infecting terrestrial isopods and nematodes. Curr Top Microbiol Immunol 116, 49–76, doi:10.1007/978-3-642-70280-8_4 (1985).

48 Poinar, G. O., Jr., Hess, R. T. & Cole, A. Replication of an Iridovirus in a nematode (Mermithidae). Intervirology 14, 316–320, doi:10.1159/000149202 (1980).

49 Widen, S. A. et al. Virus-like transposons cross the species barrier and drive the evolution of genetic incompatibilities. Science 380, eade0705, doi:10.1126/science.ade0705 (2023).

50 Widen, S. A., Bes, I. C., Koreshova, A., Krogull, D. & Burga, A., doi:10.1101/2022.07.12.499685 (2022).

51 Vance, T. D. R. & Lee, J. E. Virus and eukaryote fusogen superfamilies. Curr Biol 30, R750–R754, doi:10.1016/j.cub.2020.05.029 (2020).

52 Blanco, L. & Salas, M. Mutational analysis of bacteriophage phi 29 DNA polymerase. Methods Enzymol 262, 283–294, doi:10.1016/0076-6879(95)62024-9 (1995).

53 Blanco, L. & Salas, M. Relating structure to function in phi29 DNA polymerase. J Biol Chem 271, 8509–8512, doi:10.1074/jbc.271.15.8509 (1996).

54 Redrejo-Rodriguez, M. et al. Primer-Independent DNA Synthesis by a Family B DNA Polymerase from Self-Replicating Mobile Genetic Elements. Cell Rep 21, 1574–1587, doi:10.1016/j.celrep.2017.10.039 (2017).

55 Kazlauskas, D., Krupovic, M., Guglielmini, J., Forterre, P. & Venclovas, C. Diversity and evolution of B-family DNA polymerases. Nucleic Acids Res 48, 10142–10156, doi:10.1093/nar/gkaa760 (2020).

56 Jumper, J. et al. Highly accurate protein structure prediction with AlphaFold. Nature 596, 583–589, doi:10.1038/s41586-021-03819-2 (2021).

57 Mirdita, M. et al. ColabFold: making protein folding accessible to all. Nat Methods 19, 679–682, doi:10.1038/s41592-022-01488-1 (2022).

58 Gabler, F. et al. Protein Sequence Analysis Using the MPI Bioinformatics Toolkit. Curr Protoc Bioinformatics 72, e108, doi:10.1002/cpbi.108 (2020).

59 Shen, B. W. et al. Unusual target site disruption by the rare-cutting HNH restriction endonuclease PacI. Structure 18, 734–743, doi:10.1016/j.str.2010.03.009 (2010).

60 Stein, L. D. et al. The genome sequence of Caenorhabditis briggsae: a platform for comparative genomics. PLoS Biol 1, E45, doi:10.1371/journal.pbio.0000045 (2003).

61 Stevens, L. et al. Chromosome-Level Reference Genomes for Two Strains of Caenorhabditis briggsae: An Improved Platform for Comparative Genomics. Genome Biol Evol 14, doi:10.1093/gbe/evac042 (2022).

62 Hausner, G., Hafez, M. & Edgell, D. R. Bacterial group I introns: mobile RNA catalysts. Mob DNA 5, 8, doi:10.1186/1759-8753-5-8 (2014).

63 Lee, Y. C. et al. Single-worm long-read sequencing reveals genome diversity in free-living nematodes. Nucleic Acids Res, doi:10.1093/nar/gkad647 (2023).

64 Osman, G. A. et al. Natural Infection of C. elegans by an Oomycete Reveals a New Pathogen-Specific Immune Response. Curr Biol 28, 640–648 e645, doi:10.1016/j.cub.2018.01.029 (2018).

65 Grover, M. & Barkoulas, M. C. elegans as a new tractable host to study infections by animal pathogenic oomycetes. PLoS Pathog 17, e1009316, doi:10.1371/journal.ppat.1009316 (2021).

66 Warburton, P. E., Giordano, J., Cheung, F., Gelfand, Y. & Benson, G. Inverted repeat structure of the human genome: the X-chromosome contains a preponderance of large, highly homologous inverted repeats that contain testes genes. Genome Res 14, 1861–1869, doi:10.1101/gr.2542904 (2004).

67 Sievers, F. et al. Fast, scalable generation of high-quality protein multiple sequence alignments using Clustal Omega. Mol Syst Biol 7, 539, doi:10.1038/msb.2011.75 (2011).

68 Sievers, F. & Higgins, D. G. Clustal Omega for making accurate alignments of many protein sequences. Protein Sci 27, 135–145, doi:10.1002/pro.3290 (2018).

69 Minh, B. Q. et al. IQ-TREE 2: New Models and Efficient Methods for Phylogenetic Inference in the Genomic Era. Mol Biol Evol 37, 1530–1534, doi:10.1093/molbev/msaa015 (2020).

70 Hoang, D. T., Chernomor, O., von Haeseler, A., Minh, B. Q. & Vinh, L. S. UFBoot2: Improving the Ultrafast Bootstrap Approximation. Mol Biol Evol 35, 518–522, doi:10.1093/molbev/msx281 (2018).

71 Kalyaanamoorthy, S., Minh, B. Q., Wong, T. K. F., von Haeseler, A. & Jermiin, L. S. ModelFinder: fast model selection for accurate phylogenetic estimates. Nat Methods 14, 587–589, doi:10.1038/nmeth.4285 (2017).

72 Huerta-Cepas, J., Serra, F. & Bork, P. ETE 3: Reconstruction, Analysis, and Visualization of Phylogenomic Data. Mol Biol Evol 33, 1635–1638, doi:10.1093/molbev/msw046 (2016).

73 Larkin, M. A. et al. Clustal W and Clustal X version 2.0. Bioinformatics 23, 2947–2948, doi:10.1093/bioinformatics/btm404 (2007).

74 Tareen, A. & Kinney, J. B. Logomaker: beautiful sequence logos in Python. Bioinformatics 36, 2272–2274, doi:10.1093/bioinformatics/btz921 (2020).

75 Fu, L., Niu, B., Zhu, Z., Wu, S. & Li, W. CD-HIT: accelerated for clustering the next-generation sequencing data. Bioinformatics 28, 3150–3152, doi:10.1093/bioinformatics/bts565 (2012).

76 Capella-Gutierrez, S., Silla-Martinez, J. M. & Gabaldon, T. trimAl: a tool for automated alignment trimming in large-scale phylogenetic analyses. Bioinformatics 25, 1972–1973, doi:10.1093/bioinformatics/btp348 (2009).

77 Mirdita, M., Steinegger, M. & Soding, J. MMseqs2 desktop and local web server app for fast, interactive sequence searches. Bioinformatics 35, 2856–2858, doi:10.1093/bioinformatics/bty1057 (2019).

78 Steinegger, M. & Soding, J. MMseqs2 enables sensitive protein sequence searching for the analysis of massive data sets. Nat Biotechnol 35, 1026-1028, doi:10.1038/nbt.3988 (2017).

79 Katoh, K., Rozewicki, J. & Yamada, K. D. MAFFT online service: multiple sequence alignment, interactive sequence choice and visualization. Brief Bioinform 20, 1160–1166, doi:10.1093/bib/bbx108 (2019).

80 Ye, Y. & Godzik, A. FATCAT: a web server for flexible structure comparison and structure similarity searching. Nucleic Acids Res 32, W582–585, doi:10.1093/nar/gkh430 (2004).

81 Li, Z., Jaroszewski, L., Iyer, M., Sedova, M. & Godzik, A. FATCAT 2.0: towards a better understanding of the structural diversity of proteins. Nucleic Acids Res 48, W60–W64, doi:10.1093/nar/gkaa443 (2020).

82 Eddy, S. R. Accelerated Profile HMM Searches. PLoS Comput Biol 7, e1002195, doi:10.1371/journal.pcbi.1002195 (2011).

83 Jurka, J. Repbase update: a database and an electronic journal of repetitive elements. Trends Genet 16, 418–420, doi:10.1016/s0168-9525(00)02093-x (2000).

84 Jurka, J. et al. Repbase Update, a database of eukaryotic repetitive elements. Cytogenet Genome Res 110, 462–467, doi:10.1159/000084979 (2005).

85 Tsutakawa, S. E. et al. Crystallographic and functional studies of very short patch repair endonuclease. Mol Cell 3, 621–628, doi:10.1016/s1097-2765(00)80355-x (1999).

86 Ahmed, M. et al. Phylogenomic Analysis of the Phylum Nematoda: Conflicts and Congruences With Morphology, 18S rRNA, and Mitogenomes. Frontiers in Ecology and Evolution 9, doi:10.3389/fevo.2021.769565 (2022).

87 Tsutakawa, S. E. & Morikawa, K. The structural basis of damaged DNA recognition and endonucleolytic cleavage for very short patch repair endonuclease. Nucleic Acids Res 29, 3775–3783, doi:10.1093/nar/29.18.3775 (2001).

